# Potent neutralizing anti-SARS-CoV-2 human antibodies cure infection with SARS-CoV-2 variants in hamster model

**DOI:** 10.1101/2021.11.25.470011

**Authors:** Maya Imbrechts, Wim Maes, Louanne Ampofo, Nathalie Van den Berghe, Bas Calcoen, Dominique Van Looveren, Sam Noppen, Kevin Hollevoet, Thomas Vercruysse, Xin Zhang, Rana Abdelnabi, Caroline Foo, Hendrik Jan Thibaut, Dirk Jochmans, Karen Ven, Jeroen Lammertyn, Karen Vanhoorelbeke, Nico Callewaert, Paul De Munter, Dominique Schols, Johan Neyts, Paul Declerck, Nick Geukens

**Author notes:** corresponding author, Dr. Nick Geukens; Herestraat 49 box 820, 3000 Leuven, Belgium.

## Abstract

Treatment with neutralizing monoclonal antibodies (mAbs) against severe acute respiratory syndrome coronavirus 2 (SARS-CoV-2) contributes to COVID-19 management. Unfortunately, SARS-CoV-2 variants can escape several of these recently approved mAbs, highlighting the need for additional discovery and development. In a convalescent COVID-19 patient, we identified six mAbs, classified in four epitope groups, that potently neutralized SARS-CoV-2 Wuhan, alpha, beta, gamma and delta infection *in vitro*. In hamsters, mAbs 3E6 and 3B8 potently cured infection with SARS-CoV-2 Wuhan, beta and delta when administered post-viral infection at 5 mg/kg. Even at 0.2 mg/kg, 3B8 still reduced viral titers. Intramuscular delivery of DNA-encoded 3B8 resulted in *in vivo* mAb production of median serum levels up to 90 μg/ml, and protected hamsters against delta infection. Overall, our data mark 3B8 as a promising candidate against COVID-19, and highlight advances in both the identification and gene-based delivery of potent human mAbs.

## Introduction

The COVID-19 pandemic, caused by infection with severe acute respiratory syndrome coronavirus 2 (SARS-CoV-2), has resulted in an unprecedented global health and economic crisis and has already caused more than 5 million deaths worldwide^1^. The spike protein of SARS-CoV-2 consists of an S1 subunit that recognizes host cell receptors and an S2 subunit that promotes membrane fusion of virus and host cells. Within the S1 subunit, the receptor binding domain (RBD) is responsible for interaction with receptor angiotensin-converting enzyme 2 (ACE2) on host cells to mediate viral entry. Consequently, SARS-CoV-2 spike protein is the major target of neutralizing antibodies (Abs)^2,3^.

Antibodies can be elicited by natural infection and vaccination, or can be administered as recombinant monoclonal antibodies (mAbs) in a passive immunization strategy. Although vaccines are essential tools to fight this pandemic, therapeutic modalities, including mAbs, can also play a crucial role. This is especially the case for (immune-compromised or elderly) individuals who may not generate a robust response to their vaccine, cannot be vaccinated, are at high risk for severe illness or are still awaiting their vaccine^4^. Recent advances in mAb discovery, combined with the favorable safety profile and clinical experience, make these ideal molecules for such deployment. As of November 2021, four human mAb treatments have received (emergency use) authorization: REGN-COV2 (i.e. REGN10933 + REGN10987) from Regeneron (in the US and Europe), LY-CoV555 with/without LY-CoV016 from AbCellera/Eli Lilly (in the US), VIR-7831 from Vir Biotechnology/GlaxoSmithKline (in the US), and CT-PD9 from Celltrion (in South Korea and Europe)^5,6^.

Unfortunately, SARS-CoV-2 variants beta, gamma and delta escape from some of the currently available therapeutic mAbs. REGN10933 (i.e. one of the antibodies from the REGN-COV2 antibody cocktail) showed reduced activity against SARS-CoV-2 beta and gamma, and activity of CT-P59 was reduced against beta, gamma and delta. LY-CoV555 completely lost its activity against beta, gamma and delta, with the cocktail of LY-CoV555 and LY-CoV016 still losing activity against beta, gamma, delta plus and mu^7–13^. This demonstrates that several of the commercially available therapeutic mAbs lose their activity against the current SARS-CoV-2 variants, and that there is a need for additional potent mAbs that recognize a broad range of different SARS-CoV-2 variants.

To further broaden application and accessibility, innovations remain highly sought after in the antibody space. Gene-based delivery is one such emerging approach. Administration of the mAb sequence, using e.g. plasmid DNA (pDNA) as vector, thereby enables *in vivo* production of the mAb of interest for a prolonged period of time^14^. Compared to conventional mAb therapy, this antibody gene transfer approach can bypass the costly and complex *in vitro* protein manufacturing, facilitate combinations, and allow for a reduced administration frequency. We previously demonstrated in mice and sheep that this technology can result in *in vivo* mAb expression for several months after intramuscular pDNA delivery, in which transfection efficiency is improved by use of electroporation^15–17^. A Phase I trial of a DNA-encoded mAb against Zika virus was initiated in 2019 (NTC: NCT03831503, sponsor: Inovio Pharmaceuticals), further illustrating the advances in clinical translation. In the context of COVID-19, various funding organizations, including the US Defense Advanced Research Projects Agency (DARPA), have dedicated considerable funding to the development of gene-based delivery of SARS-CoV-2 neutralizing mAbs. Indeed, progress in mAb discovery and innovative delivery technologies can revolutionize emerging infectious diseases responses.

In this work, we sought to identify human mAbs reactive to all current SARS-CoV-2 variants both *in vitro* and *in vivo*, and explore antibody gene transfer. We were able to generate highly potent and broadly neutralizing mAbs that are capable of treating SARS-CoV-2 Wuhan, beta and delta infection in Syrian Golden hamsters. We demonstrated efficacy both as recombinant protein and encoded in pDNA, highlighting innovation in human mAb discovery and gene-based delivery.

## Results

### 1. Identification and characterization of anti-SARS-CoV-2 antibodies from convalescent COVID-19 patients

To identify fully human SARS-CoV-2 neutralizing antibodies, we first analyzed RBD-binding titers and neutralizing antibody titers in serum, as well as the percentage RBD-positive B cells in PBMCs from 26 convalescent COVID-19 patients (see Supplementary Table 1). Patient K-COV19-901, having both a high titer of neutralizing antibodies and a high number of RBD-specific B cells, was selected for the isolation of individual RBD-specific B cells via fluorescence-activated cell sorting (FACS), using biotinylated RBD (Wuhan isolate) combined with streptavidin-PE as bait (Supplementary Figure 1). Next, these single B cells were used as starting material for our single B cell cloning strategy, where we amplified the coding sequence of the IgG antibody heavy and light chain variable domains and combined them with the human IgG1 constant domain in a vector for recombinant mAb production. A panel of 20 unique mAb sequences (labelled K-COV-901-X, further mentioned as “X”) was selected for *in vitro* production and subsequent characterization. All 20 antibodies retained RBD and trimeric spike antigen (Wuhan) binding *in vitro* when analyzed via ELISA or surface plasmon resonance (SPR) assays. In addition, antibodies showed picomolar affinities to the RBD antigen and trimeric spike antigen with equilibrium constants (K_D_ values) ranging from 31 to 443 pM and 32 to 243 pM, respectively (Table 1).

**Table 1:** Overview of antigen binding, affinity and neutralizing capacity of 20 selected antibodies.

| mAb | ELISA antigen binding | | Affinity $K_D$ (pM) | | Neutralization $IC_{50}$ (nM) |
| --- | --- | --- | --- | --- | --- |
|  | RBD (Wuhan) | Trimeric spike (Wuhan) | RBD (Wuhan) | Trimeric spike (Wuhan) | Trimeric spike (D614G) |
| **1A10** | + | + | **199 ± 58** | **89 ± 45** | **0.83** |
| 1B11 | + | + | 443 ± 4 | 89 ± 18 | / |
| **1C1** | + | + | **41 ± 12** | **44 ± 12** | **0.78** |
| **1C11** | + | + | **87 ± 24** | **84 ± 29** | **0.20** |
| 1D5 | + | + | 111 ± 13 | 127 ± 78 | 15.74 |
| 1B5 | + | + | 45 ± 2 | 56 ± 16 | 1.83 |
| 2A2 | + | + | 220 ± 17 | 176 ± 88 | 27.53 |
| 2A8 | + | + | 78 ± 8 | 74 ± 5 | 8.91 |
| **2B11** | + | + | **48 ± 21** | **32 ± 8** | **0.71** |
| **2B2** | + | + | **31 ± 7** | **63 ± 14** | **0.18** |
| 2C8 | + | + | 64 ± 8 | 91 ± 15 | / |
| 2D10 | + | + | 52 ± 8 | 65 ± 36 | 36.28 |
| **2D6** | + | + | **31 ± 6** | **69 ± 18** | **0.51** |
| 3A11 | + | + | 58 ± 8 | 91 ± 18 | 21.67 |
| 3B3 | + | + | 252 ± 21 | 99 ± 18 | / |
| **3B8** | + | + | **234 ± 104** | **86 ± 12** | **0.02** |
| 3C11 | + | + | 48 ± 10 | 71 ± 11 | 12.69 |
| **3E6** | + | + | **373 ± 295** | **243 ± 163** | **0.35** |
| 3E9 | + | + | 248 ± 13 | 143 ± 20 | 1.09 |
| 3F9 | + | + | 98 ± 16 | 141 ± 168 | / |

Antigen binding was evaluated for RBD and trimeric spike protein (Wuhan) via ELISA (in duplicate) and SPR (min. triplicate). Affinity ( $K_D$ ; in pM; average ± standard deviation of at least 3 replicates) was determined via SPR. Neutralizing capacity is given in 50% inhibitory concentration ( $IC_{50}$ ; in nM) and was measured in triplicate via a SARS-CoV-2 D614G VSV pseudovirus assay. Bold and italic = 8 best neutralizing antibodies; + = potent antigen binding ( $OD \geq 2$ ) at mAb concentration of 1 μg/ml or lower; / = no neutralizing activity when tested at a concentration of 5 μg/ml or 33.34 nM.

Next, we evaluated the functional activity of these antibodies *in vitro.* To this end, we used a pseudovirus assay with vesicular stomatitis virus (VSV) expressing the SARS-CoV-2 spike protein (D614G) on its surface and evaluated pseudovirus infection of Vero E6 cells in the presence and absence of each of the antibodies. Out of 20 mAbs, 16 could neutralize SARS-CoV-2 pseudovirus infection, with 8 very potent mAbs having a half-maximal 50% inhibitory concentration (IC_50_) lower than 1 nM and mAb 3B8 having an IC_50_ of only 0.017 nM (Figure 1 & Table 1). Given their potent neutralizing capacity, these 8 mAbs were selected for further characterization.

**Figure 1:**
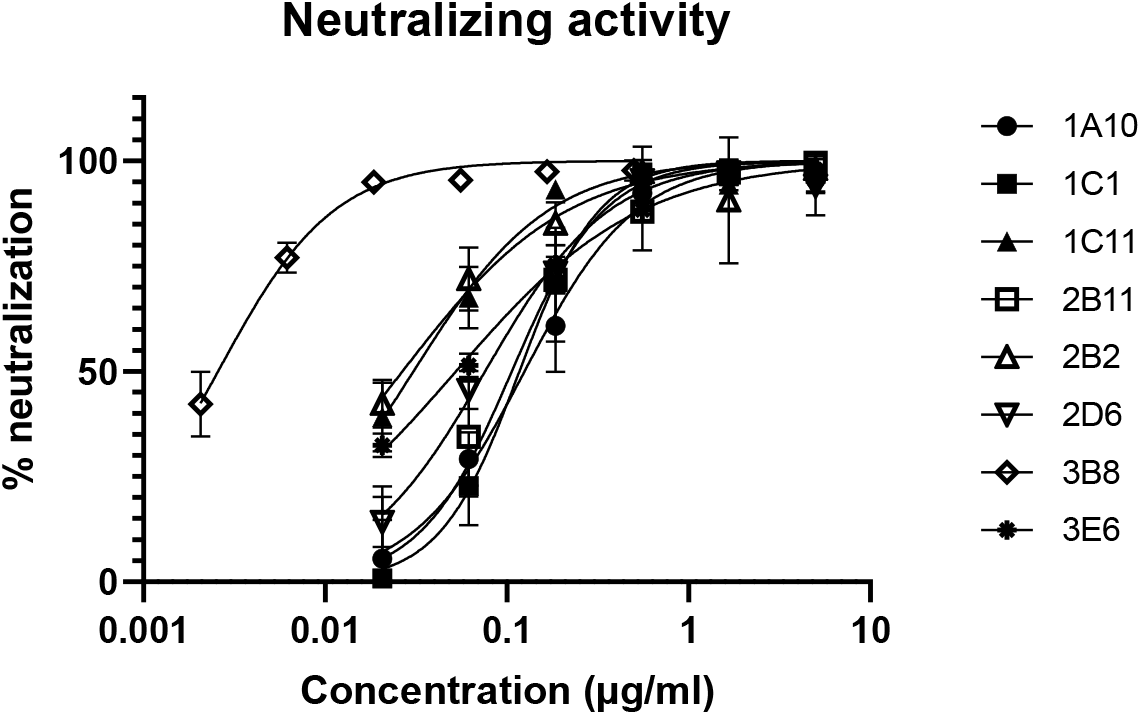
Neutralizing activity of top 8 most potent antibodies. Percentage neutralizing activity, measured via a SARS-CoV-2 D614G VSV pseudotyped assay, is given at different concentrations for the top 8 antibodies. Symbols show mean and standard deviation of 3 replicates, graph is representative for 2 independent experiments.

### 2. Evaluation of *in vitro* efficacy against SARS-CoV-2 variants and other coronavirus species

With the continuous emergence of new SARS-CoV-2 variants, it was crucial to evaluate antigen binding and neutralizing capacity of the selected antibodies against these variants as well. RBD antigens bearing the single mutations E484K, E484Q, N501Y and L452R, present – alone or combined – in multiple SARS-CoV-2 variants (i.e. alpha, beta, gamma, delta and others), as well as RBD antigens bearing the double mutation E484Q, L452R or triple mutation K417N, E484K, N501Y, as present in SARS-CoV-2 variants kappa and beta respectively^18^, were used to assess binding and affinity of the 8 most potent antibodies via both ELISA and SPR. In addition, RBD mutations N439K, described as an antibody evasion mutant^19–21^, and RBD Y453F, originating from Danish mink farms and a potential neutralization resistance mutation^22,23^, were evaluated (an overview of the analyzed RBD single mutations and their presence in SARS-CoV-2 variants is given in Supplementary Table 2). Interestingly, 7 out of 8 antibodies retained their binding capacity against all analyzed RBD single mutant antigens. One mAb, 2B11, showed reduced binding to RBD L452R and did not bind RBD E484K. This mAb did neither bind the antigens with multiple mutations that included L452R or E484K. mAb 1C11, although binding with all single mutant antigens, lost its activity against RBD L452R, E484Q double mutant antigen (Table 2 top part). All antibodies that bound the single mutant RBD antigens, still showed high affinities to these antigens, as evident from the equilibrium constants (K_D_) ranging from 9 – 647 pM.

**Table 2:**
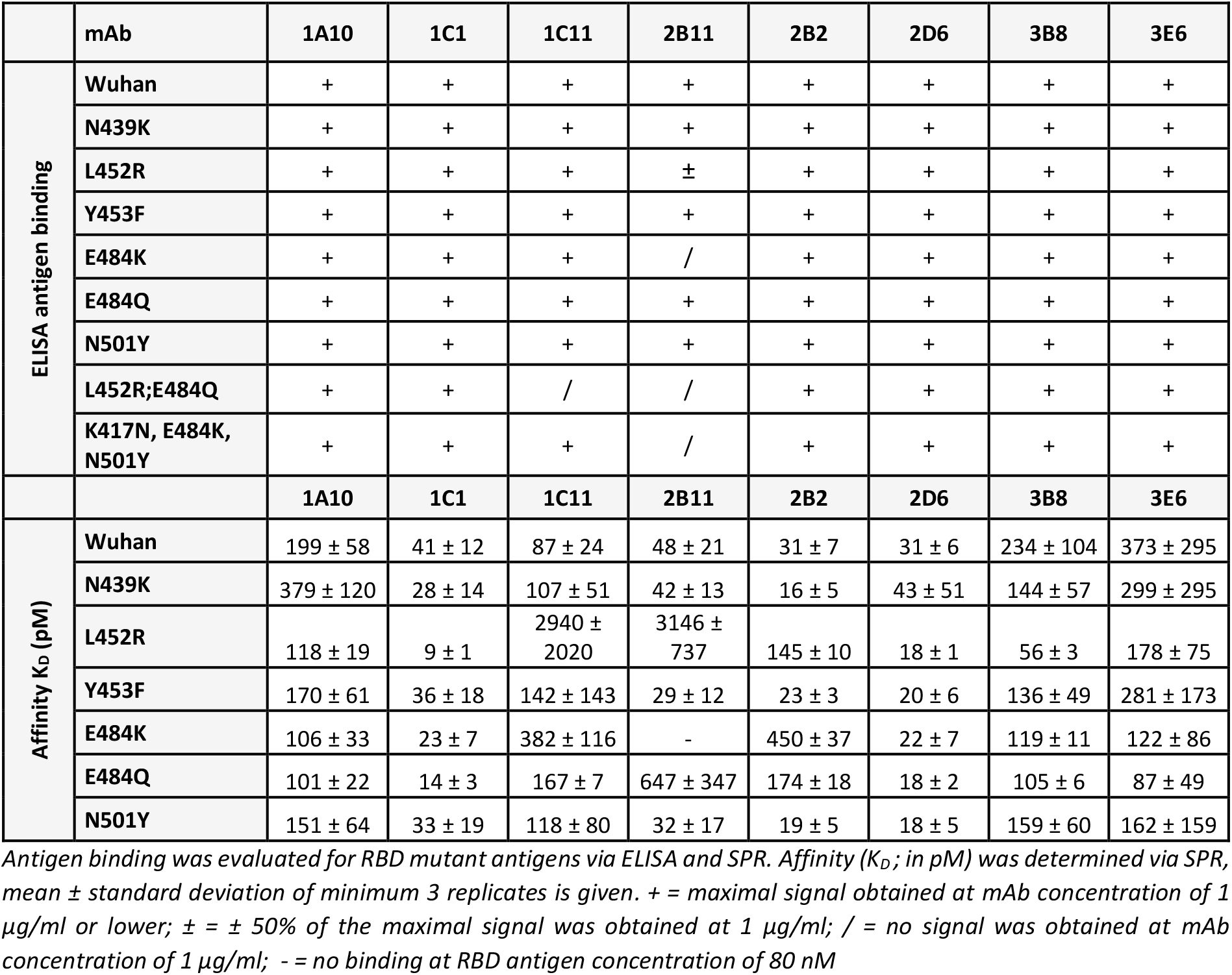
Overview of binding and affinity of top 8 mAbs with RBD mutant antigens.

|  | mAb | 1A10 | 1C1 | 1C11 | 2B11 | 2B2 | 2D6 | 3B8 | 3E6 |
| --- | --- | --- | --- | --- | --- | --- | --- | --- | --- |
| ELISA antigen binding | Wuhan | + | + | + | + | + | + | + | + |
|  | N439K | + | + | + | + | + | + | + | + |
|  | L452R | + | + | + | ± | + | + | + | + |
|  | Y453F | + | + | + | + | + | + | + | + |
|  | E484K | + | + | + | / | + | + | + | + |
|  | E484Q | + | + | + | + | + | + | + | + |
|  | N501Y | + | + | + | + | + | + | + | + |
|  | L452R;E484Q | + | + | / | / | + | + | + | + |
|  | K417N, E484K, N501Y | + | + | + | / | + | + | + | + |
|  |  | 1A10 | 1C1 | 1C11 | 2B11 | 2B2 | 2D6 | 3B8 | 3E6 |
| Affinity $K_D$ (pM) | Wuhan | 199 ± 58 | 41 ± 12 | 87 ± 24 | 48 ± 21 | 31 ± 7 | 31 ± 6 | 234 ± 104 | 373 ± 295 |
|  | N439K | 379 ± 120 | 28 ± 14 | 107 ± 51 | 42 ± 13 | 16 ± 5 | 43 ± 51 | 144 ± 57 | 299 ± 295 |
|  | L452R | 118 ± 19 | 9 ± 1 | 2940 ± 2020 | 3146 ± 737 | 145 ± 10 | 18 ± 1 | 56 ± 3 | 178 ± 75 |
|  | Y453F | 170 ± 61 | 36 ± 18 | 142 ± 143 | 29 ± 12 | 23 ± 3 | 20 ± 6 | 136 ± 49 | 281 ± 173 |
|  | E484K | 106 ± 33 | 23 ± 7 | 382 ± 116 | - | 450 ± 37 | 22 ± 7 | 119 ± 11 | 122 ± 86 |
|  | E484Q | 101 ± 22 | 14 ± 3 | 167 ± 7 | 647 ± 347 | 174 ± 18 | 18 ± 2 | 105 ± 6 | 87 ± 49 |
|  | N501Y | 151 ± 64 | 33 ± 19 | 118 ± 80 | 32 ± 17 | 19 ± 5 | 18 ± 5 | 159 ± 60 | 162 ± 159 |

Antigen binding was evaluated for RBD mutant antigens via ELISA and SPR. Affinity ( $K_D$ ; in pM) was determined via SPR, mean ± standard deviation of minimum 3 replicates is given. + = maximal signal obtained at mAb concentration of 1 µg/ml or lower; ± = ± 50% of the maximal signal was obtained at 1 µg/ml; / = no signal was obtained at mAb concentration of 1 µg/ml; - = no binding at RBD antigen concentration of 80 nM

Only antibodies 1C11 and 2B11 showed an increased K_D_ value for RBD L452R (2940 pM and 3146 pM respectively) (Table 2 bottom part).

In addition to antigen binding, functional activity against SARS-CoV-2 variants was evaluated in a cytopathic effect neutralization test (CPENT) using live SARS-CoV-2 Wuhan, alpha, beta and gamma virus strains and Vero E6 cells, as well as in a plaque reduction neutralization test (PRNT) using live SARS-CoV-2 delta strain with the same cell line. Infection with SARS-CoV-2 Wuhan and alpha was best neutralized by antibodies 3B8 and 2B2 (IC_50_ ≤ 0.63 and ≤ 0.16 nM respectively), while SARS-CoV-2 beta infection was best neutralized by antibodies 3B8 and 3E6 (IC_50_ ≤ 0.32 nM). The same was true for SARS-CoV-2 gamma (IC_50_ ≤ 0.37 nM for 3B8 and 3E6), although the high standard deviations hinder drawing strong conclusions. SARS-CoV-2 delta was most potently neutralized by 3B8 (IC_50_ = 0.01 nM), with next in line antibodies 1C1, 2D6, 3E6 (IC_50_ values between 0.98 – 1.08 nM). On the other hand, antibodies 2B11 and 1C11 lost their neutralizing activity against SARS-CoV-2 delta (IC_50_ > 67 nM). Overall, mAb 3B8 was the most potent neutralizing mAb towards all variants, with IC_50_ values ranging from 0.37 nM to 0.01 nM for neutralization of SARS-CoV-2 gamma and delta strain respectively (Figure 2 and Table 3).

**Figure 2:**
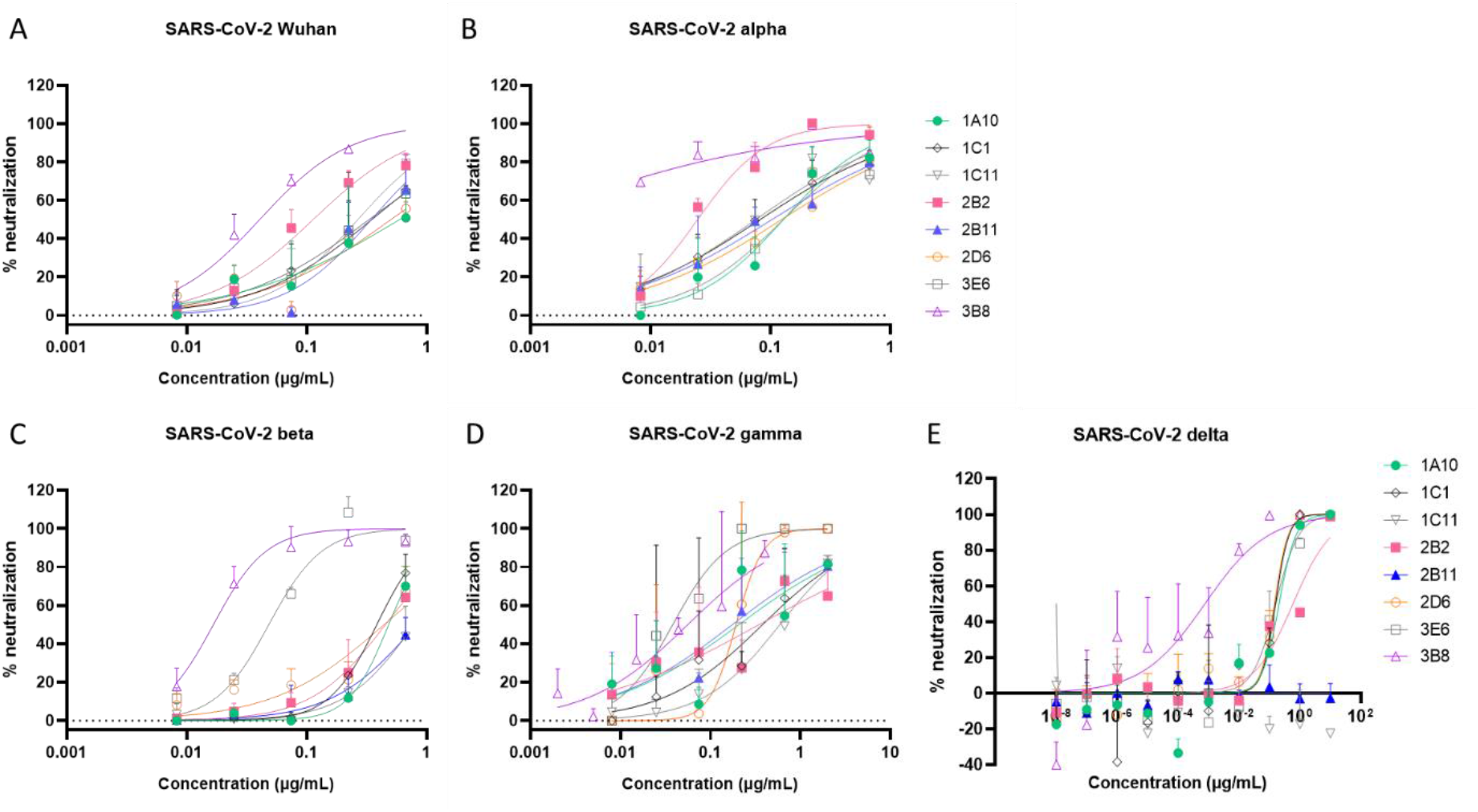
*in vitro* neutralization of infection with live SARS-CoV-2 variants. Concentration-dependent neutralizing activity (% neutralization) was measured via CPENT assay using SARS-CoV-2 Wuhan, alpha, beta and gamma (A-D) or via PRNT assay using SARS-CoV-2 delta (E) for the 8 selected antibodies. Symbols show mean and standard deviation (only upper bar is shown for clarity) of 3 (A-D) or 2 (E) replicates.

**Table 3:** Overview of *in vitro* neutralizing activity (IC_50_) towards SARS-CoV-2 Wuhan, alpha, beta, gamma and delta for top 8 mAbs.

| mAb | Neutralization - $IC_{50}$ (nM) | | | | |
| --- | --- | --- | --- | --- | --- |
|  | CPE live virus assay – Wuhan | CPE live virus assay – alpha | CPE live virus assay – beta | CPE live virus assay – gamma | PRNT live virus assay – delta |
| 1A10 | 3.94 | 0.85 | 3.21 | 1.38 | 1.39 |
| 1C1 | 2.31 | 0.55 | 2.56 | 2.53 | 1.05 |
| 1C11 | 1.53 | 0.9 | 5.23 | 3.92 | ND |
| 2B11 | 2.29 | 0.69 | 5.46 | 1.17 | ND |
| 2B2 | 0.63 | 0.16 | 3.02 | 2.31 | 3.97 |
| 2D6 | 3.34 | 0.82 | 2.94 | 1.28 | 0.98 |
| 3B8 | 0.25 | 0.1 | 0.11 | 0.37 | 0.01 |
| 3E6 | 2.19 | 0.85 | 0.32 | 0.19 | 1.08 |

Neutralizing activity ( $IC_{50}$ ; in nM) was determined via CPE assay for SARS-CoV-2 Wuhan, alpha, beta and gamma and via PRNT assay for SARS-CoV-2 delta.

The eight selected neutralizing antibodies were also tested for their ability to cross-neutralize other human coronavirus strains using VSV pseudotypes expressing the spike protein from SARS-CoV, Middle East respiratory stress (MERS)-CoV or 229E-CoV on their surface. However, none of the selected antibodies was able to cross-neutralize one of these coronavirus species when used at a concentration of 16.67 nM (or 2.5 μg/ml) (data not shown).

### 3. Classification of mAb epitopes

The selected mAbs were classified in different categories based on their epitope specificity using an ELISA cross-competition assay where the antibodies were pairwise tested against one another. Two groups could be distinguished consisting of antibodies 1A10, 1C1 and 2D6 on the one hand and antibodies 2B2, 2B11 and 1C11 on the other hand. Antibodies 3B8 and 3E6 could not be classified in one of these groups (Figure 3). This suggests that the 8 selected mAbs bind to 4 different regions of the RBD antigen.

**Figure 3:**
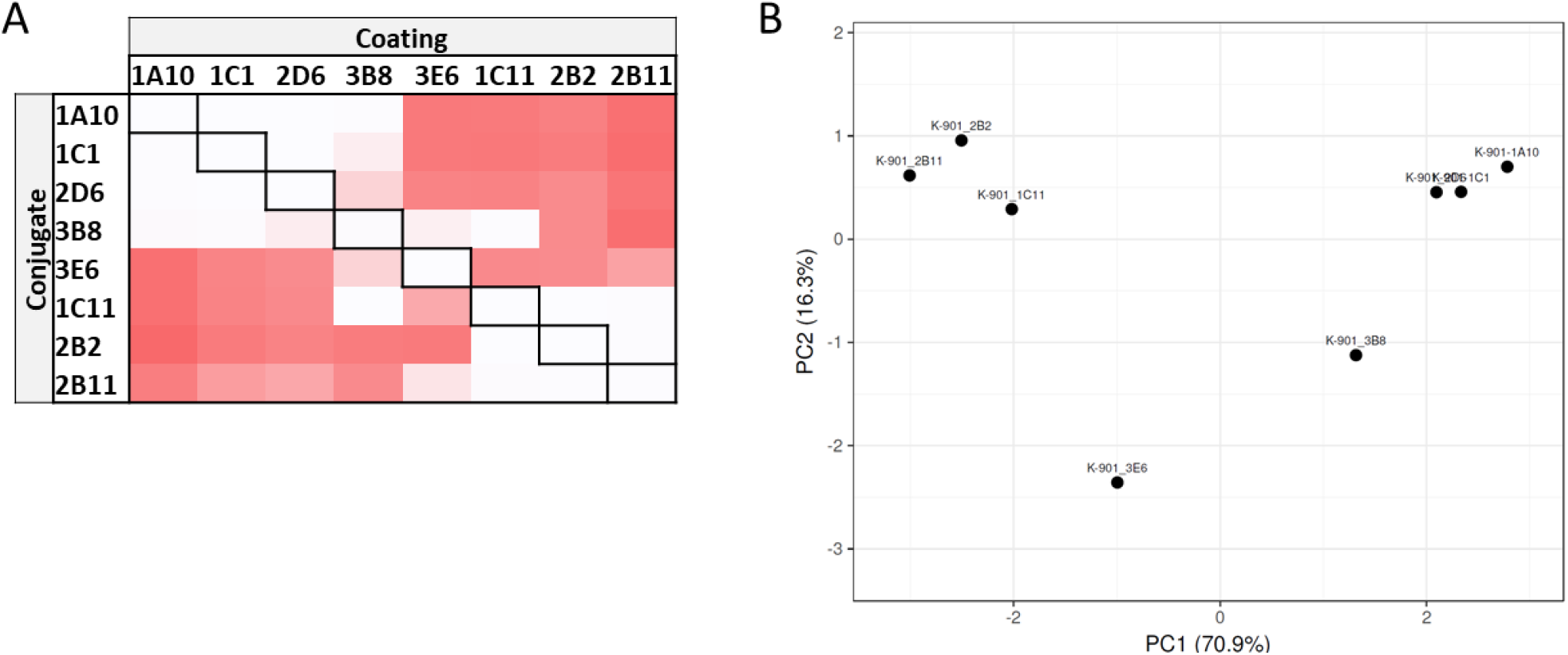
Epitope specificity of top 8 mAbs. A) Visual representation of the cross-competition ELISA. All antibodies were pairwise tested against one another as coating or detection mAb. Signal strength is visualized by color intensity, with a white color indicating no binding of the detection mAb could occur, and dark red indicating strong binding of the detection mAb. B) Principal component analysis of the results obtained in the cross-competition ELISA to enable classification of the antibodies in different epitope groups. Data are representative for 2 independent experiments.

### 4. Potent *in vivo* efficacy on different SARS-CoV-2 variants in the Syrian golden hamster model

Antibodies 3B8, 3E6, 2B2 and 1C1 were selected for *in vivo* evaluation based on their *in vitro* neutralizing activity combined with their variability in epitope binding. In a first study, 30 hamsters were divided into 5 groups (6 animals/group) and infected intranasally with 50 μl of SARS-CoV-2 Wuhan virus suspension (containing approximately 2×10^6^ tissue culture infective dose (TCID_50_)). After 24h, antibodies 3B8, 3E6, 2B2, 1C1 or a human IgG1 isotype control were injected intraperitoneally at a dose of 5 mg/kg. All animals were sacrificed at 4 days post infection for analysis of viral RNA and viral infectious titer in the lungs, as well as mAb concentration in serum (Figure 4a). The presence of therapeutic RBD-specific human antibodies circulating in the blood of the treated hamsters was verified through ELISA. This revealed that 5 animals (1 out of 6 for clone 3E6 and 2 out of 6 for clones 2B2 and 1C1) were not successfully injected (Figure 4b). We therefore excluded all animals without detectable serum levels from further analysis. In all groups, mAb treatment reduced viral RNA load in lung tissue as compared to the isotype control group with statistical significance (p < 0.05 for 2B2; p < 0.01 for 3B8, 3E6, 1C1) (Figure 4c). No infectious virus in the lung tissue could be detected in any of the animals treated with mAb 3B8. In animals treated with antibodies 3E6 and 2B2, a low infectious titer was detected in 1 animal, while three out of four animals treated with mAb 1C1 had a low detectable titer. In all groups, the difference was statistically significant (p < 0.01) compared to the isotype control group, thus showing good *in vivo* efficacy for the treatment of SARS-CoV-2 (Wuhan) infection (Figure 4d).

**Figure 4:**
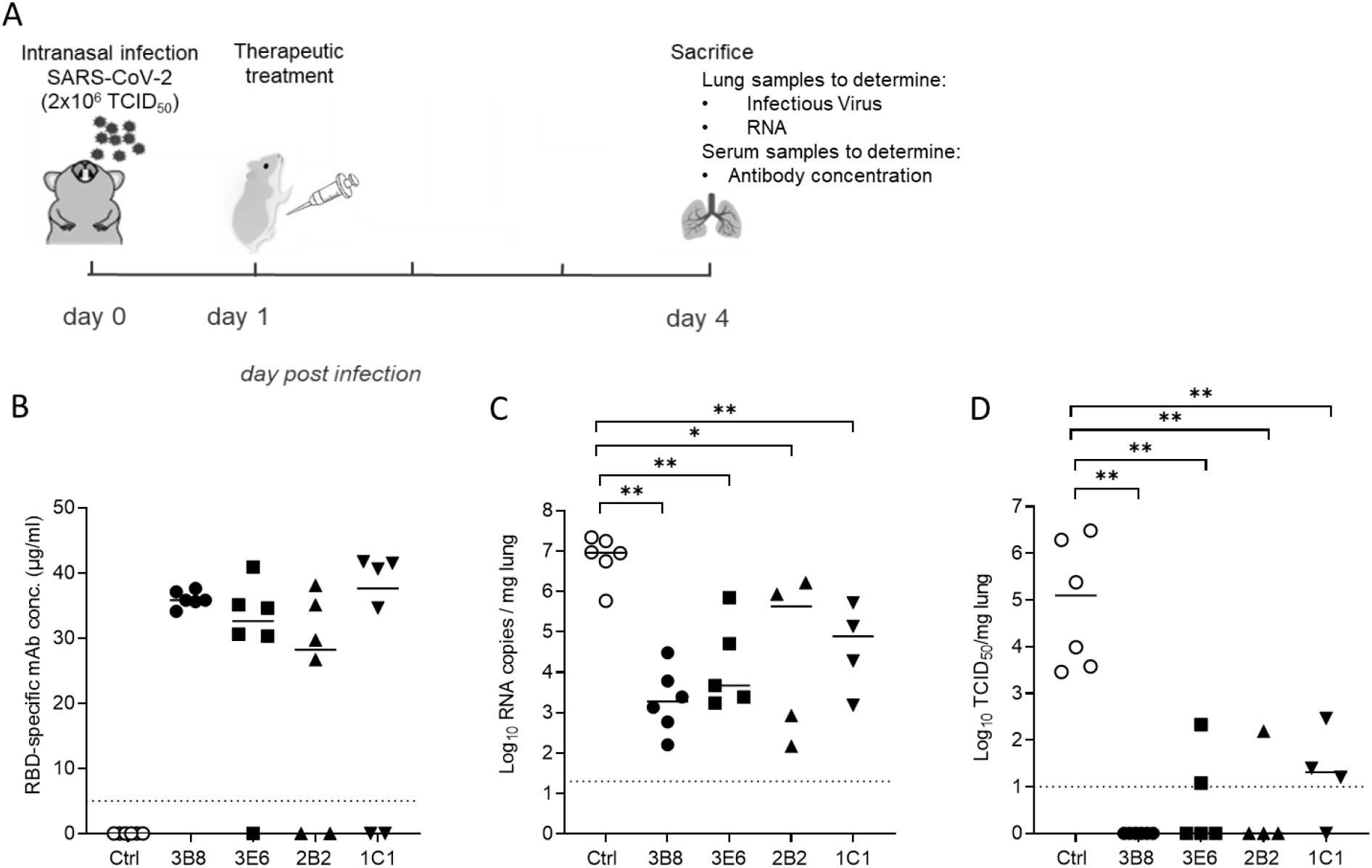
Treatment of SARS-CoV-2 Wuhan infection by antibodies 3B8, 3E6, 2B2 and 1C1 (5 mg/kg) in hamsters. A) Study design. Hamsters were intranasally infected with SARS-CoV-2 on day 0 and received intraperitoneal mAb treatment (5 mg/kg) 24h post infection. Animals were sacrificed at day 4 for analysis of lung and blood samples. B) Concentration (μg/ml) of the administered antibodies in serum at day 4 post infection (n=6 per group). Animals without detectable mAb serum levels were excluded from the graphs in panel C and D. C) Viral RNA levels, expressed as log_10_ RNA copies per mg of lung tissue, on day 4 post infection. D) Infectious viral loads, expressed as log_10_ TCID_50_ per mg lung tissue, at day 4 post infection. Individual data with median values are shown. Dotted line represents the detection limit. * = p < 0.05; ** = p < 0.01 as determined via Mann-Whitney U test.

As it was shown previously that SARS-CoV-2 variants of concern beta and delta are more resistant to neutralization by vaccine-induced antibodies or currently available mAb treatments^7,24,25^, and given that SARS-CoV-2 delta is currently the most dominant variant worldwide^26,27^, we decided to evaluate *in vivo* treatment efficacy of our antibodies against both variants as well. A study design similar to the first study was used, with 30 hamsters divided in 5 groups and intranasally infected with 50 μl of SARS-CoV-2 beta^28^ or SARS-CoV-2 delta suspension (both containing approximately 1×10^4^ TCID_50_). Animals were injected intraperitoneally with IgG1 isotype control or antibodies 3B8, 3E6, 2B2 and REGN-COV-2 at 5 mg/kg 24h post infection and were sacrificed 4 days post infection, with REGN-COV-2 antibody cocktail included for benchmarking purposes^29^. In the study using SARS-CoV-2 beta, one animal from groups 3B8, 2B2, 1C1 and REGN-CoV-2 as well as two animals from group 3E6 were excluded from analysis because of the absence of detectable antibody serum levels (Figure 5a). In addition, it should be noted that two animals, one from group 3B8 and one from group 2B2, showed very low antibody serum titers, but were not excluded from analysis (indicated by open symbols in the respective groups). Viral RNA load in lung tissue was significantly reduced by treatment with mAb 3B8, 3E6 and REGN-CoV-2 compared to isotype control (p < 0.05 for 3B8 and 3E6; p < 0.01 for REGN-CoV-2), while the reduction observed for 2B2 was not statistically significant (Figure 5b). No infectious viral titer in lung tissue (TCID_50_) could be detected upon treatment with 3B8, 3E6 or REGN-CoV-2 (p < 0.01 compared to isotype control), while a viral titer was still detectable in four out of five animals treated with 2B2 (Figure 5c). When analyzing the efficacy of our antibodies against SARS-CoV-2 delta infection, a similar picture was observed. Here, only one animal from group 2B2 was excluded because of a lack of detectable serum concentrations (Figure 5d). All treatments resulted in a significant reduction (p < 0.01) of both viral load and infectious viral titers in lung tissue (Figure 5 e and f), with no detectable viral titers in animals treated with antibodies 3B8, 3E6 and REGN-CoV-2. In conclusion, treatment with antibodies 3B8 and 3E6 at 24h post infection resulted in a statistically significant reduction of virus in lung tissue of animals infected with SARS-CoV-2 beta or delta. Although significantly reducing infectious viral titers in lung tissue of animals infected with SARS-CoV-2 delta, mAb 2B2 seems to be less potent as treatment for SARS-CoV-2 beta and delta infection compared to antibodies 3B8 and 3E6, an observation corresponding to the *in vitro* neutralization data for these variants.

**Figure 5:**
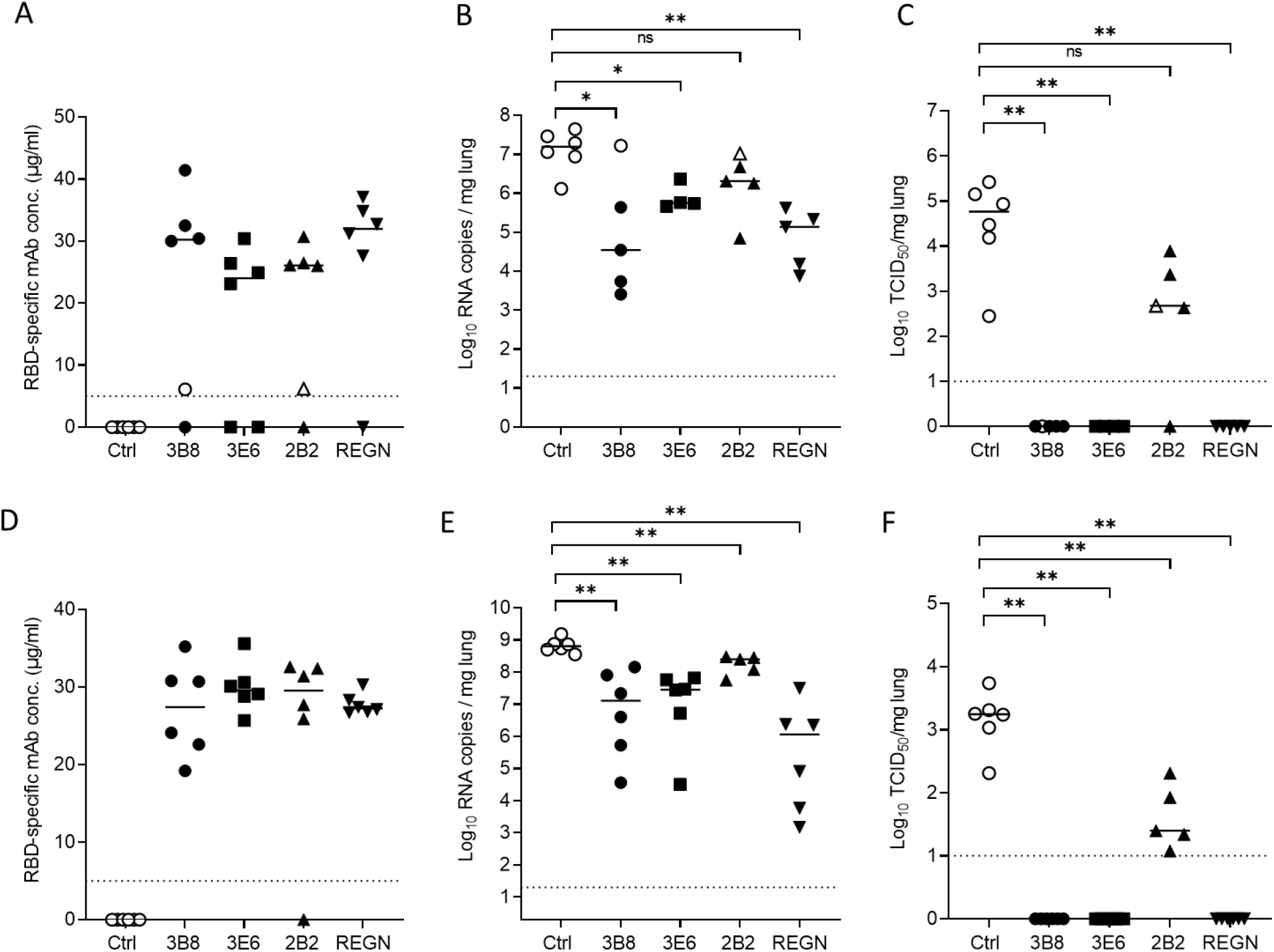
Treatment of SARS-CoV-2 beta and delta infection by antibodies 3B8, 3E6, 2B2 and REGN-COV-2 (5 mg/kg) in hamsters. Concentration (μg/ml) of the administered antibodies at day 4 post infection in serum from hamsters (n=6 per group) infected with SARS-CoV-2 beta (A) or delta (D). Viral RNA levels in the lungs of hamsters infected with SARS-CoV-2 beta (B) or delta (E), on day 4 post infection. Infectious viral loads, in lungs of hamsters infected with SARS-CoV-2 beta (C) or delta (F), on day 4 post infection. Animals without detectable antibody serum levels were excluded from the graphs in panel B-C-E-F. Individual data and median values are shown. Dotted line represents the detection limit. ns = p > 0.05; * = p < 0.05; ** = p < 0.01 as determined via Mann-Whitney U test.

### 5. Dose-dependent therapeutic efficacy of 3B8

Antibodies 3B8 and 3E6 both completely abrogated infectious viral titers after infection with SARS-CoV-2 Wuhan, beta and delta when administered 24h post infection at 5 mg/kg. As 3B8 was superior to 3E6 in *in vitro* neutralization experiments, this mAb was used in an *in vivo* dose-response experiment. Hamsters were intranasally infected with SARS-CoV-2 delta (1×10^4^ TCID_50_). At 24h post infection, isotype control (5 mg/kg) or mAb 3B8 (5 mg/kg; 1 mg/kg, 0.2 mg/kg or 0.04 mg/kg) was administered intraperitoneally. Animals were sacrificed at day 4 post infection. No animals were excluded from analysis based on a lack of detectable antibody serum levels, although one animal from group 0.2 mg/kg had serum levels 3 times lower compared to its group members (animal indicated by open symbol) (Figure 6a). Treatment with a dose of 5 mg/kg and 1 mg/kg of 3B8 significantly reduced the amount of viral RNA detected in lung tissue, and resulted in undetectable levels of infectious virus. 5, 1 and 0.2 mg/kg showed significant differences in viral RNA (p < 0.05 or < 0.01 for 5 vs 1 mg/kg or 5 vs 0.2/0.04 mg/kg respectively; p < 0.01 for 1 vs 0.2 or 0.04 mg/kg), indicating a positive dose-response effect. When dosing at 0.2 mg/kg, no statistically significant (p=0.13) decrease in viral RNA was seen, but infectious virus was undetectable in lung tissue of 3 out of 6 animals, and was decreased (more than 4 times lower than lower limit of 95% confidence interval from isotype control group) in a fourth animal. In the animal having reduced antibody titers (open symbol), no reduction in viral titer could be observed. Also, at the lowest dose of 0.04 mg/kg, no effect could be observed on viral RNA or infectious viral titer (Figure 6b and c).

**Figure 6:**
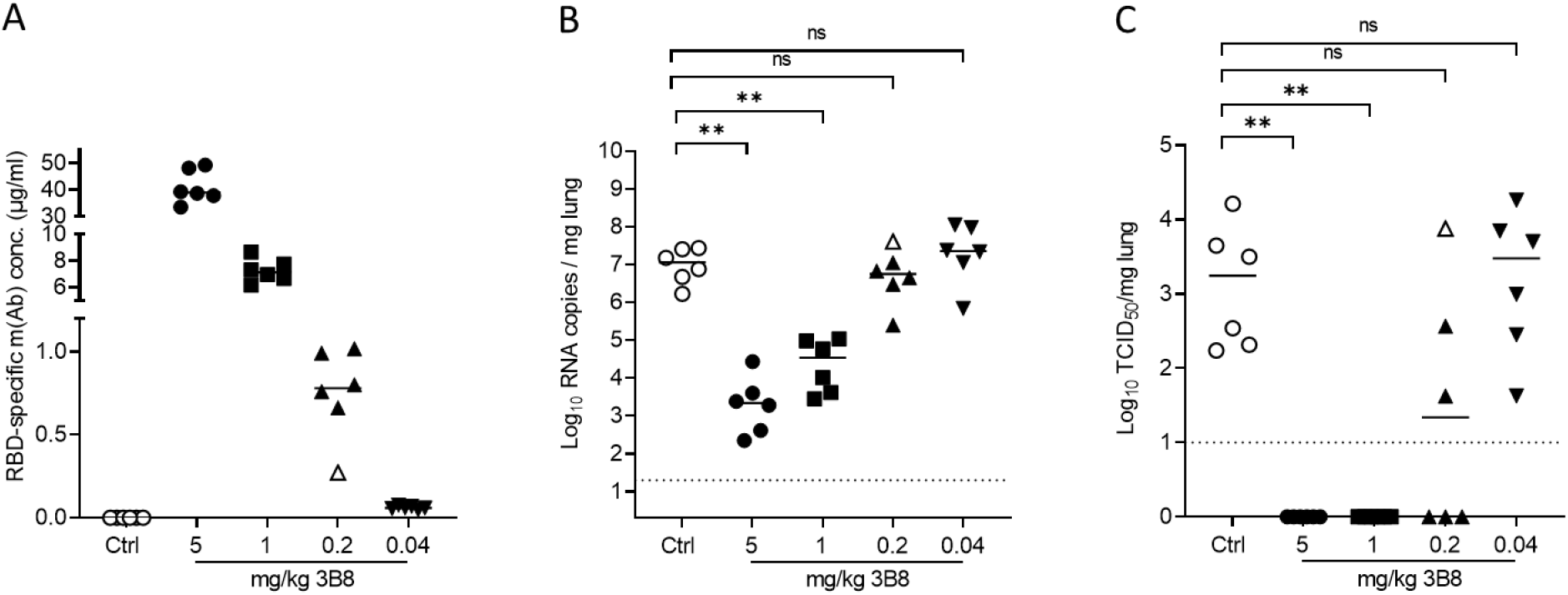
Dose-response experiment for treatment of SARS-CoV-2 delta infection with mAb 3B8 (5, 1, 0.2 or 0.04 mg/kg) in hamsters. A) Concentration (μg/ml) of the administered antibodies in serum at day 4 post infection (n=6 per group). B) Viral RNA levels, expressed as log_10_ RNA copies per mg of lung tissue, on day 4 post infection. C) Infectious viral loads, expressed as log_10_ TCID_50_ per mg lung tissue, at day 4 post infection. Individual data and median values are shown. Dotted line represents the detection limit. ns = p > 0.05; ** = p < 0.01 as determined via Mann-Whitney U test.

### 6. Intramuscular delivery of DNA-based 3B8

To assess whether 3B8 could be produced *in vivo* at sufficient titers to protect from SARS-CoV-2 delta infection, intramuscular electroporation of DNA-encoded 3B8 (p3B8) was performed in hamsters either on day 10, day 7 or day 5 prior to infection. Median 3B8 serum levels at day 0, the day of intranasal infection, ranged between 30 μg/ml (p3B8 at day -5) and 90 μg/ml (p3B8 at day -10) (Figure 7A). All transfected hamsters displayed detectable mAb titers. The longer the lag time between p3B8 delivery and infection, the higher the resulting serum 3B8 concentrations at day 0. This was anticipated, as *in vivo* expressed mAb levels typically increase and accumulate in the first two weeks after intramuscular pDNA administration^15,17^. Of note, mAb titers did not consistently or significantly increase between day 0 and day 4 post infection, and a markedly higher variability was observed at day 4 (Figure 7A). These observations are likely linked to the interaction of 3B8 with the virus (including target-mediated clearance) from day 0 on. Irrespective of the timing of pDNA administration, the *in vivo* produced 3B8 levels were sufficient to protect the animals from viral challenge. Compared to the untreated control group, all animals showed a significantly lower amount of viral RNA in lung tissue, and undetectable levels of infectious virus (except for one animal at day -7) (Figure 7B-C). No 3B8 dose-response effect was observed, since all timepoints gave a profound protection. These data show how intramuscular injection of DNA-based 3B8 provides a robust *in vivo* mAb expression and protection against SARS-CoV-2 delta infection.

**Figure 7:**
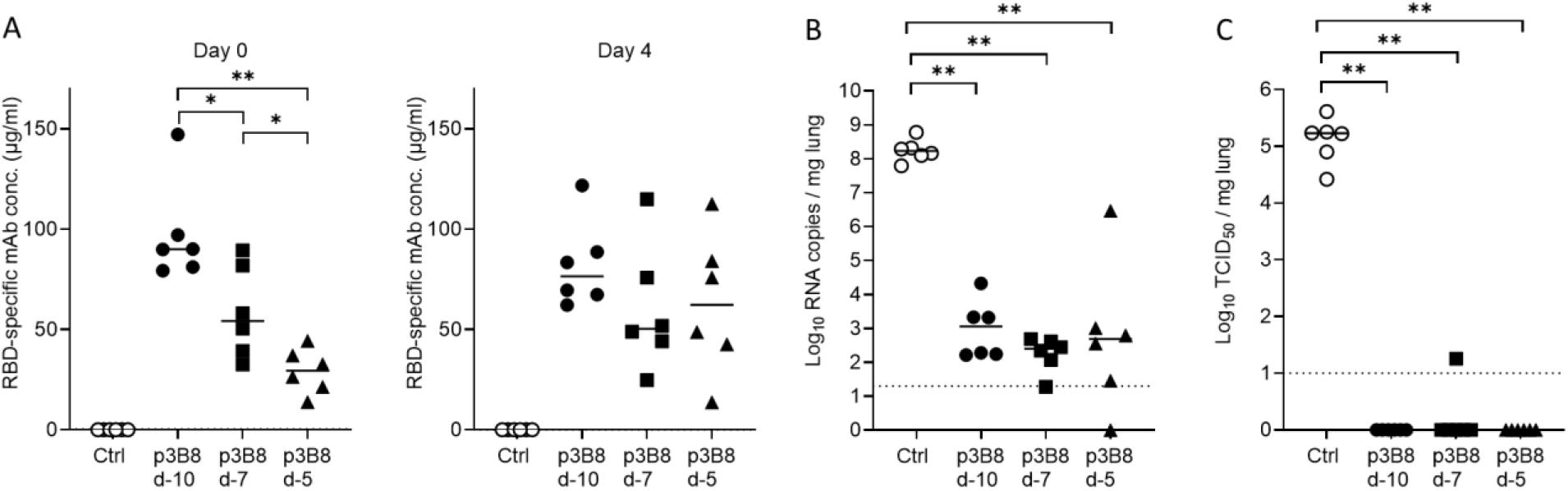
Pharmacokinetics and efficacy of intramuscular DNA-encoded 3B8 delivery for protection against SARS-CoV-2 delta infection in hamsters. 600 μg p3B8 was delivered via intramuscular electroporation at either 10, 7 or 5 days prior to infection (n=6 per group). Animals of the negative control group were left untreated. A) Concentration (μg/ml) of 3B8 in serum at day 0 and 4 post infection. B) Viral RNA levels, expressed as log_10_ RNA copies per mg of lung tissue, on day 4 post infection. C) Infectious viral loads, expressed as log_10_ TCID_50_ per mg lung tissue, at day 4 post infection. Individual data and median values are shown. Dotted line represents the detection limit. ** = p < 0.01 as determined via Mann-Whitney U test.

## Discussion

Despite vaccination rates steadily increasing, the majority of individuals worldwide have not been fully vaccinated and the number of SARS-CoV-2 related hospitalizations and deaths continue to increase^1^. In addition, vaccine-breakthrough infections are becoming more prevalent. These may be explained by waning vaccine protection, and the emergence of SARS-CoV-2 variants with reduced sensitivity to vaccine-elicited antibody neutralization^7,8,38–41,30–37^. mAb treatments can therefore play a crucial role in addressing COVID-19 morbidity and mortality and to prevent rapid dissemination of viral infection in specific settings, such as elderly care homes.

In this study, eight highly potent neutralizing mAbs (IC_50_ < 1 nM) were identified from a convalescent COVID-19 patient, by leveraging our B cell mining platform. Six out of eight antibodies bound all analyzed recombinant SARS-CoV-2 RBD antigens bearing single or multiple mutations. Affinities were evaluated for RBD antigens with single mutations and shown to be in the picomolar range. Corresponding to the antigen binding data, *in vitro* neutralization assays using live SARS-CoV-2 Wuhan, alpha, beta, gamma and delta strains showed that these antibodies potently cross-neutralized all variants with IC_50_ values ranging from 0.01 to 3.97 nM. Interestingly, data from the cross-competition ELISA showed that these antibodies were not all targeting the same epitope, but could be divided into four epitope groups.

The most potent mAb of each of these epitope classes was tested for therapeutic activity *in vivo* in Syrian golden hamsters. This model has been widely used to assess the efficacy of vaccines and therapeutics against SARS-CoV-2 infection as it closely mimics the clinical disease observed in humans^42–44^. Using this model, we designed three different therapeutic studies to evaluate the efficacy of our antibodies as a treatment for SARS-CoV-2 Wuhan, beta and delta infection. We showed that antibodies 3B8 and 3E6 reduced median viral RNA titers with a factor 50 - 5000 or 10 - 330 respectively when used as a treatment (5 mg/kg) for all tested SARS-CoV-2 strains. In addition, treatment with these antibodies resulted in undetectable levels of infectious virus in the lungs of almost all animals. Our *in vitro* and *in vivo* data suggest that 3B8 and 3E6 target an epitope conserved between the different variants, abolishing the need for combination therapy for the current SARS-CoV-2 variants. In addition, since we identified potent neutralizing antibodies from different epitope groups, combination therapy remains an option in the future if additional variants of concern appear. Indeed, thanks to the advancements in mAb identification and characterization at display in this study, we can rapidly identify multiple mAbs of varying specificity and potency, which in combination could result in enhanced breadth and potency.

Importantly, we also showed that 3B8 is an ultrapotent therapeutic mAb, with treatment at a dose of 1 mg/kg resulting in undetectable infectious viral titers. Even when administered at 0.2 mg/kg, viral replication was reduced, with median infectious viral titers of the treatment group being 67 times lower compared to the isotype control group. This further highlights the potency of mAb 3B8 in a therapeutic setting^2,44–48^. High potencies are crucial to decrease cost of goods, enable sustainable manufacturability and increase the number of doses produced annually, and may therefore also facilitate the availability of therapeutic mAbs to low- and middle-income countries.

When comparing the mAbs identified in this study with the commercially available ones, we noticed that K_D_ values observed for our antibodies were consistently between 3- and 200-fold lower than values reported for marketed antibodies of Vir Biotechnology, Regeneron Pharmaceuticals or Eli Lilly^49–51^. However, when analyzing K_D_ values of REGN-COV2 antibodies 10933 and 10987 in our own experiments for benchmarking purposes, K_D_ values were similar to values observed for our mAbs (data not shown). In addition, it should be noted that antibodies 3B8 and 3E6 each performed equally well as monotherapy (when administered at 5 mg/kg) to treat SARS-CoV-2 infection beta and delta in hamsters when compared to the marketed REGN-COV2 antibody cocktail, which consists of two different antibodies^29^. Moreover, we could observe therapeutic efficacy of mAb 3B8 *in vivo* at a dose of only 0.2 mg/kg. Interestingly, to the best of our knowledge, no *in vivo* therapeutic mAb efficacy data have been published yet for SARS-CoV-2 delta infection in hamsters. In addition, only for REGN-COV2 some therapeutic efficacy was reported in hamsters at such low mAb doses, although after infection of hamsters with SARS-CoV-2 Wuhan and using different readouts^42^.

In addition to demonstrating our ability to select and characterize highly potent neutralizing human mAbs from a COVID-19 patient, we evaluated the DNA-based delivery of 3B8, an innovative delivery approach that could revolutionize the application of mAb therapeutics for infectious diseases^14,52^. We found that intramuscular electroporation of p3B8 provided a robust *in vivo* mAb expression and protection against SARS-CoV-2 delta infection, irrespective of the timepoint of p3B8 administration. The resulting mAb serum concentrations produced *in vivo* were similar to or significantly higher than what was observed after i.p. injection of 5 mg/kg 3B8 protein. As a consequence, lower pDNA doses and/or alternative timepoints (closer to or after viral infection), are anticipated to still demonstrate efficacy. To the best of our knowledge, this is the first report to show that *in vivo* DNA-based delivery of a SARS-Cov-2-neutralizing mAb can protect from viral infection. Such approach can complement and accelerate the discovery, development and delivery of (combinations of) mAbs, and allow us to keep up the pace with the current and future pandemics.

Although no clear evidence for antibody-dependent disease enhancement (ADE) in COVID-19 patients has been reported until now, it has been suggested that its potential risk should be continuously monitored^4,9,44,53^. In this light, it should be emphasized that the high potency of 3B8 is obtained without the contribution of Fc-mediated effector functions, as fully human IgG1 antibodies were not species-matched before administration to hamsters. This suggests that mAb 3B8 will tolerate the future introduction of Fc-effector silence mutations to avoid any potential risk of ADE, favoring its further evaluation as a future mAb treatment.

In conclusion, the high therapeutic potency of mAb 3B8 *in vivo* against SARS-CoV-2 delta, achieved both as protein and encoded in plasmid, combined with its broad cross-reactivity against all other tested SARS-CoV-2 variants, make 3B8 a very interesting candidate to help fight the COVID-19 pandemic. The combination of accelerated human mAb discovery and innovative gene-based delivery, at display in the current study, holds the potential to revolutionize emerging infectious diseases responses.

## Materials & methods

### 1. Isolation of PBMCs from convalescent COVID-19 patients

Patients with a PCR-confirmed SARS-CoV-2 infection (>18 years old; recovered from disease) were recruited from the University Hospital Leuven or AZ Groeninge Kortrijk for peripheral blood collection. The study and corresponding experiments were approved by the local ethics committee (S64089) and all patients gave their written informed consent. Immediately after blood sample collection, peripheral blood mononuclear cells (PBMCs) were isolated from EDTA-treated blood by density centrifugation (Lymphoprep, STEMCELL Technologies Inc). After washing, the collected PBMCs were resuspended in freezing medium consisting of 10% (v/v) dimethyl sulfoxide (DMSO; Merck) and 90% fetal bovine serum (FBS; GIBCO) for cryopreservation at −80°C or in liquid nitrogen for long term storage.

### 2. Single-cell sorting of antigen-specific B cells

Human B cells were enriched from cryopreserved PBMCs using the EasySep^™^ Human B Cell Enrichment Kit (STEMCELL Technologies) according to the manufacturer’s instructions. After purification, B cells were stained with Zombie Aqua^™^ Fixable Viability Kit (Biolegend) and blocked with FcR Blocking Reagent (Miltenyi Biotec) for 15 minutes on ice. Following the 15 minutes incubation, B cells were washed with FACS buffer (phosphate-buffered saline (PBS) + 2% FBS + 2 mM EDTA) and incubated with biotinylated SARS-CoV-2 RBD protein for 60 minutes at on ice. His-tag labeled SARS-CoV-2 RBD (The Native Antigen Company) was biotinylated with the EZ-Link Sulfo-NHS-LC-Biotin kit (Thermofisher Scientific) according to the manufacturer’s protocol, corresponding to 1-3 biotin groups per antibody molecule. After the 60 minutes incubation, B cells were washed with FACS buffer and stained with PerCP-cy5.5 anti-human CD19 antibody (Biolegend, 363016), FITC anti-human CD3 antibody (Biolegend, 300306) and PE streptavidin (Biolegend, 405203) for 25 minutes on ice. Subsequently, the stained cells were washed twice with FACS buffer and single-cell sorted by BD Influx^™^ Cell Sorter (BD Biosciences). UltraComp eBeads^™^ Compensation Beads and tosylactivated M-280 Dynabeads (Invitrogen) coupled with SARS-CoV-2 RBD protein according to the manufacturer’s instructions were used for compensation. Selection of RBD-specific B cells was performed using the following gating strategy: lymphocytes were selected using FSC and SSC, followed by selection of single cells based on width versus area of FSC and SSC signals. Next, live cells were selected as Zombia Aqua negative cells. Here, B cells were selected as CD19+ CD3-cells and evaluated for RBD surface staining. To allow proper gating of RBD-positive cells, a fluorescence minus one (FMO) control was included (supplementary figure 1). Individual B cells were sorted into 96-wells PCR plates (Bioké) containing 2 μL lysis buffer (0.2% (v/v) Triton X-100 + 2U/μL RNaseOUT^™^ Recombinant Ribonuclease (Invitrogen) in UltraPure^™^ DNAse/RNase free distilled water (Invitrogen)) per well. The plates were sealed with a Microseal^®^ ‘F’ Film (BioRad) and immediately frozen on dry ice before storage at −80°C for further use.

### 3. Amplification of antibody variable domains

Transcripts of lysed single B cells were denatured and hybridized with a mix of 1 μL 10 mM each nucleotide dNTP-Mix (Invitrogen) and 1 μL 10 μM oligo-dT primer (5’-AAGCAGTGGTATCAACGCAGAGTACT_30_-3’) at 72°C for 3 minutes. Subsequently, cDNA synthesis and pre-amplification was performed according to Picelli et al^54^. cDNA was stored at −20°C. cDNA was purified with Agencourt AMPure XP beads (Beckman Coulter) according to the manufacturer’s protocol and quantified on the BioAnalyzer 2100 instrument (Agilent) and on a Qubit^™^ fluorometer (Thermofisher Scientific). The sequences encoding the immunoglobulin (Ig) heavy- and light-chain variable regions (V_H_ and V_L_) of IgM and IgG were amplified using reverse primers described by Ozawa et al^55^ and forward primer described by Picelli et al^54^. The PCR reaction was performed with KAPA HiFi HotStart ReadyMix (Roche) at 95°C for 3 min, followed by 30 cycles at 98°C for 20 seconds, 60°C for 45 seconds and 72°C for 60 seconds and terminated at 72°C for 5 minutes. All PCR products were checked on a 1% agarose gel and purified using the Agencourt AMPure XP beads. Concentrations were measured on a Qubit^™^ fluorometer for Sanger sequencing (outsourced).

### 4. Recombinant antibody production and purification

#### a. Generation and amplification of expression vectors containing the selected heavy- and light chain sequences

Generation of antibody expression vectors was outsourced (Genscript). The Ig V_H_ and V_L_ sequences were cloned into a pcDNA3.4 expression vector containing human IgG1 constant regions. The plasmid DNA (pDNA) was amplified by transformation in DH5α *E. coli* and subsequent purification using the NucleoBond Xtra Maxi EF kit (Machery - Nagel) according to the manufacturer’s instructions. Plasmid purity and integrity was checked on a 1% agarose gel.

#### b. Production of recombinant antibody in HEK93F cells

Recombinant antibodies were transiently transfected and produced in vitro in 293F Freestyle suspension cells (Thermofisher Scientific). Briefly, equal amounts of heavy- and light chain pDNA were mixed with X-tremeGENE HP DNA Transfection Reagent (Roche) in FreeStyle^™^ 293 Expression Medium (Thermofisher Scientific) according to the manufacturer’s protocol and incubated for at least 15 minutes to allow the pDNA to enter the liposomes. The transfection reagent mixture was added in a dropwise manner to 293F Freestyle suspension cells. The cells were cultured in T175 flasks (Sarstedt, Germany) on an orbital shaker (Thermofischer Scientific) at 135 rpm and 8% CO_2_ in a 37°C humidified incubator for 5 days. At day 5, supernatant containing the recombinant antibodies was collected by centrifugation (1,400 x g for 10 minutes at room temperature), filtered through a 0.2 μm filter unit (VWR) and stored at −20°C.

#### c. Purification of recombinant antibody

Purification of the recombinant antibodies was conducted on a ÄKTAprime plus system (GE Healthcare Life Sciences) at 4°C using a 1 mL HiTrap^®^ Protein A HP pre-packed Protein A Sepharose^®^ column (Cytiva). Briefly, the column was first equilibrated in buffer A (20 mM sodium phosphate, 150mM NaCl, pH 7.5) before the sample was loaded on the column. The flow rate for all steps was 1 mL/min. The column was washed with buffer A and buffer B (20 mM Sodium Phosphate, 500 mM NaCl, pH 7.5). The recombinant antibodies were eluted from the column with 100 mM sodium acetate, pH 3.5 and the collected fractions were immediately neutralized with 1M Tris, pH 9. The fractions were pooled and dialyzed against sterile-filtered PBS. The ÄKTAprime plus system and all glasswork was treated with 30% hydrogen peroxide (VWR) to reduce the presence of endotoxin.

#### d. Quality control using SDS-PAGE and endotoxin measurements

Purified recombinant antibodies were assessed by SDS-PAGE under nonreducing conditions using the Amersham Phastsystem^™^ (GE Healthcare) according to the manufacturer’s instructions. Endotoxin levels were measured on Endosafe^®^-PTS^™^ (Charles River) according to the manufacturer’s instructions.

### 5. ELISA

#### a. Antigen binding

The binding of the purified recombinant antibodies to the following SARS-CoV-2 antigens was assessed via ELISA: spike glycoprotein (S1) RBD-His (REC31849-500, The Native Antigen Company), RBD(N439K)-His (40592-V08H14, Sino Biological), RBD(N501Y)-His (40592-V08H82, Sino Biological), RBD(E484K)-His (40592-V08H84, Sino Biological), RBD(Y453F)-His (40592-V08H80, Sino Biological), RBD(E484Q)-His (40592-V08H81, Sino Biological), RBD(L425R)-His (40592-V08H28, Sino Biological), RBD(L425R,E484Q)-His (40592-V08H88, Sino Biological), RBD(K417N, E484K, N501Y)-His (40592-V08H85, Sino Biological), Spike Glycoprotein (Full-Length)-His (REC31868-500, The Native Antigen Company). Briefly, polystyrene 96-well flat-bottom microtiter plates (Corning costar) were coated with 100 μL/well of 2 μg/mL antigen diluted in PBS and incubated overnight at 4°C. The next day, plates were blocked with 200 μL/well blocking buffer containing 1% (w/v) Bovine Serum Albumin (BSA, Sigma Aldrich) in PBS for 2 hours at room temperature. The plates were then washed six times with wash buffer (0.002% (v/v) Tween-80 (Sigma Aldrich) in PBS). After washing, 100 μL/well of the samples, calibrators and/or controls were added. As a positive and negative control, an anti-SARS-CoV-2 RBD mAb (40150-D004, Sino Biological) and anti-SARS-CoV-2 nucleocapsid antibody (MBS2563841, MyBioSource) were used, respectively. All samples and controls were diluted in PBS + 0.1% (w/v) BSA + 0.002% (v/v) Tween 80 with or without 1,86 g/L EDTA (PTA(E) buffer). After incubation, the plates were washed six times and incubated with 100 μL/well horseradish peroxidase (HRP)-conjugated goat anti-human IgG (Fc-specific, Sigma Aldrich) diluted 1/5000 in PTA buffer. After 1 hour incubation at room temperature, the plate was again washed six times and 100 μL/well substrate (200 μL 40 mg/mL o-Phenylenediamine dihydrochloride 99+% (OPD, Acros Organics BVBA) + 2 μL H_2_O_2_ (Merck) in 20 mL citrate buffer, pH 5 (0.1 M citric acid monohydrate from Sigma and 0.2 M disodium phosphate dihydrate from Merck)) was added to the plate and incubated for 30 minutes in the dark. The color reaction was stopped with 50 μL/well 4 M H_2_SO_4_ (Thermofisher Scientific). Optical density (OD) was measured at 492 nm with an ELx808 ELISA reader (BioTek). Analysis was performed using Graphpad Prism 9.0 (Graphpad Software).

#### b. Competition assay

Polystyrene 96-well flat-bottom microtiter plates were coated with 4 μg/mL purified recombinant capture antibody in PBS (100 μL/well) and incubated at 4°C. After an overnight incubation, plates were blocked with 200 μL/well blocking buffer for 2 hours at room temperature. The plates were washed six times with washing buffer and 10 ng/mL SARS-CoV-2 RBD protein diluted in PTA buffer (100 μL/well) was added to the plates for a 2-hour incubation at room temperature. After the incubation, the plates were washed and captured antigen was detected with 100 μL/well of 1/100 biotin-conjugated purified recombinant antibodies diluted in PTA buffer. Biotinylation of the purified recombinant mAbs was performed with the EZ-Link Sulfo-NHS-LC-Biotin kit according to the manufacturer’s protocol. After a 1-hour incubation at room temperature and washing, biotinylated recombinant antibodies that were able to bind the antigen were detected with 1/10,000 poly-HRP-conjugated streptavidin (Sanquin) diluted in PTA buffer (100 μL/well) and incubated for an additional 30 minutes at room temperature. After a final washing step, 100 μL/well substrate was added to the plate and incubated at room temperature for 30 minutes in the dark at room. The color reaction was stopped with 50 μL/well 4 M H_2_SO_4_. Optical density (OD) was measured at 492 nm with an ELx808 ELISA reader (BioTek). Very low OD values indicate the detection antibody was not able to bind, suggesting similar epitopes of both coating and detection antibodies, while high OD values indicate both antibodies have a different epitope. Epitope binning graphs were made in Microsoft Excel and clustering was done with ClustVis web tool (https://biit.cs.ut.ee/clustvis/).

### 6. Surface plasmon resonance

Surface Plasmon Resonance (SPR) was used to evaluate the interaction between monoclonal Abs and SARS-CoV-2 antigens (i.e. full-length spike protein, RBD and RBD mutants; see catalog numbers under Methods section “ELISA”). The binding experiments were performed at 25°C on a Biacore T200 instrument (GE Healthcare, Uppsala, Sweden) in HBS-EP+ buffer (10 mM HEPES, 150 mM NaCl, 3 mM EDTA and 0.05% v/v Surfactant P20). First, mouse anti-human IgG (Fc) antibody (Human Antibody Capture Kit, Cytiva) was immobilized on a CM5 chip according to manufacturer instructions. Monoclonal Abs were then captured between 30 and 100 RU. Increasing concentrations of analyte were sequentially injected in one single cycle at a flow rate of 30 μl/min. The dissociation was monitored for 30 min. The chip was finally regenerated with 3M MgCl2 before a new mAb was captured. A reference flow was used as a control for non-specific binding and refractive index changes. Several buffer blanks were used for double referencing. Binding affinities (KD) were derived after fitting the experimental data to the 1:1 binding model in the Biacore T200 Evaluation Software 3.1 using the single cycle kinetic procedure. Each interaction was repeated a least three times.

### 7. Monoclonal antibody neutralization assays

#### a. Production of S-pseudotyped virus and serum neutralization test (SARS2, SARS1, MERS, 229E)

VSV S-pseudotypes were generated as described previously^56^. Briefly, HEK-293T cells (SARS-CoV, SARS-CoV-2 and MERS-S) or BHK-21J cells (229E) were transfected with the respective S protein expression plasmids, and one day later infected (MOI = 2) with GFP-encoding VSVΔG backbone virus (purchased from Kerafast). Two hours later, the medium was replaced by medium containing anti-VSV-G antibody (I1-hybridoma, ATCC CRL-2700) to neutralize residual VSV-G input. After 24 h incubation at 32 °C, the supernatants were harvested. To quantify nAbs, serial dilutions of serum samples were incubated for 1 hour at 37 °C with an equal volume of S pseudotyped VSV particles and inoculated on Vero E6 cells (SARS-CoV and SARS-CoV-2) or Huh-7 cells (229E and MERS-S) for 18 hours. The percentage of GFP expressing cells was quantified on a Cell Insight CX5/7 High Content Screening platform (Thermo Fischer Scientific) with Thermo Fisher Scientific HCS Studio (v.6.6.0) software. Neutralization IC_50_ values were determined by normalizing the serum neutralization dilution curve to a virus (100%) and cell control (0%) and fitting in Graphpad Prism (GraphPad Software, Inc.).

#### b. Cell-based cytopathic (CPE) assay and CPE-based virus neutralization test (CPENT) (SARS-CoV-2 Wuhan, alpha, beta, gamma)

The titers of the virus stocks (TCID50) and 50% neutralizing titers (log_10_CPENT_50_) were determined using cytopathic effect (CPE)-based cell assays and CPE-based virus neutralization tests (CPENT), respectively, on VeroE6 cells with some modifications. Shortly, 10^4^ VeroE6 cells/well were seeded in 96-well plates one day prior to the titration and inoculated with 10–fold serial dilutions of virus solutions and cultured for 3 days at 37°C. Later, assays first visually scored for CPE and then stained with MTS/Phenazine methosulphate (PMS; Sigma-Aldrich) solution for 1.5 h at 37 °C in the dark. Post MTS/PMS staining absorbance was measured at 498 nm for each well. All assays were performed in six replicates and TCID_50_/ml was determined using Reed and Muench method^57^. CPENT was measured in a similar way as the CPE assays for viral titration with some modifications. In brief, serial dilutions of antibodies were mixed separately with live SARS-CoV-2 Wuhan, alpha, beta and gamma virus strains, incubated at 37 °C for 1h, and added to the monolayer of Vero E6 cells. All distinct antibodies were assayed in triplicate in serial dilutions. CPE neutralization was calculated with the following formula: % neutralization activity = % CPE reduction = (OD _Virus + antibody_ – OD _VC_) / (OD _CC_ – OD _VC_) *100 and 50% neutralization titers (CPENT_50_) were calculated with non-linear regression. Cells either unexposed to the virus or mixed with TCID_50_ SARS-CoV-2 were used as negative (uninfected) and positive (infected) controls, respectively.

#### c. SARS-CoV-2 plaque reduction neutralization test (PRNT) (SARS-CoV-2 delta)

The PRNT experiments were performed with a SARS-CoV-2 isolate from the B.1.617.2 lineage. This virus was isolated from oropharyngeal swabs taken from a patient in Belgium (EPI_ISL_2425097; 2021-04-20). Virus stocks were prepared by 2x passaging on Vero E6 cells.

Dose-dependent neutralization of distinct antibodies was assessed by mixing the antibodies at different concentrations, with 100 PFU SARS-CoV-2 in DMEM supplemented with 2% FBS and incubating the mixture at 37°C for 1h. Antibody-virus complexes were then added to VeroE6 cell monolayers in 12-well plates and incubated at 37°C for 1h. Subsequently, the inoculum mixture was replaced with 0.8% (w/v) methylcellulose in DMEM supplemented with 2% FBS. After 3 days incubation at 37°C, the overlays were removed, the cells were fixed with 3.7% PFA, and stained with 0.5% crystal violet. Half-maximum neutralization titers (PRNT50) were defined as the antibody concentration that resulted in a plaque reduction of 50%.

### 8. Golden Syrian hamster studies

#### a. SARS-CoV-2

Three SARS-CoV-2 strains were used in this study. BetaCov/Belgium/GHB-03021/2020 (EPI ISL 109 40797612020-02-03), was recovered from a nasopharyngeal swab taken from an RT-qPCR confirmed asymptomatic patient who returned from Wuhan, China in the beginning of February 2020. A close relation with the prototypic Wuhan-Hu-1 2019-nCoV (GenBank accession 112 number MN908947.3) strain was confirmed by phylogenetic analysis. Infectious virus was isolated by serial passaging on Huh7 and Vero E6 cells^58^; passage 6 virus was used for the study described here. The Beta variant B.1.351 (hCoV-19/Belgium/rega-1920/2021; EPI_ISL_896474, 2021-01-11) was isolated from nasopharyngeal swabs taken from a traveler returning to Belgium in January 2021 who became a patient with respiratory symptoms. The Delta variant, B.1.617.2 (hCoV-19/Belgium/rega-7214/2021; EPI_ISL_2425097; 2021-04-20) was isolated from nasopharyngeal swabs in April 2021 in Belgium through active surveillance. Both strains were subjected to sequencing on a MinION platform (Oxford pore) directly from the nasopharyngeal swabs. Infectious virus was isolated by passaging on VeroE6 cells^28^; passage 2 was used for the study described here. Live virus-related work was conducted in the high-containment A3 and BSL3+ facilities of the KU Leuven Rega Institute (3CAPS) under licenses AMV 30112018 SBB 219 2018 0892 and AMV 23102017 SBB 219 20170589 according to institutional guidelines.

#### b. Cells

Vero E6 cells (African green monkey kidney, ATCC CRL-1586) were cultured in minimal essential medium (MEM, Gibco) supplemented with 10% fetal bovine serum (Integro), 1% non-essential amino acids (NEAA, Gibco), 1% L-glutamine (Gibco) and 1% bicarbonate (Gibco). End-point titrations on Vero E6 cells were performed with medium containing 2% fetal bovine serum instead of 10%. Huh7 cells were cultured in Dulbecco’s modified Eagle’s medium (DMEM, Gibco) supplemented with 10% fetal bovine serum (Integro), 1% bicarbonate (Gibco), 2% HEPES buffer (Gibco). Both cells were kept in a humidified 5% CO_2_ incubator at 37°C.

#### c. SARS-CoV-2 infection model in hamsters

The hamster infection model of SARS-CoV-2 has been described before^58,59^. Female Syrian hamsters *(Mesocricetus auratus)* were purchased from Janvier Laboratories and kept per two in individually ventilated isolator cages (IsoCage N Bio-containment System, Tecniplast) at 21°C, 55% humidity and 12:12 day/night cycles. Housing conditions and experimental procedures were approved by the ethics committee of animal experimentation of KU Leuven (license P065-2020). For infection, female hamsters of 6-8 weeks old and 75-100 g were anesthetized with ketamine/xylazine/atropine and inoculated intranasally with 50 μL containing either 2×10^6^ TCID50 SARS-CoV-2 (ancestral Wuhan strain), 1×10^4^ TCID50 (Beta B.1.351), or 1×10^4^ TCID50 (Delta B.1.617.2). Antibody protein treatments (anti-SARS-CoV-2 mAbs or human IgG1 isotype control Trastuzumab/Herceptin^®^ (Roche)) were initiated 24 hours post infection by intraperitoneal injection. Intramuscular pDNA electroporation was done at d-10, d-7 and d-5 infection. Hamsters were monitored for appearance, behavior, and weight.

At day 4 post-infection, animals were euthanized by intraperitoneal injection of 500 μL Dolethal (200 mg/mL sodium pentobarbital, Vétoquinol SA). Lungs were collected and viral RNA and infectious virus were quantified by RT-qPCR and end-point virus titration, respectively. Blood samples were collected at end-point for pharmacokinetic analysis. No randomization methods were used and confounders were not controlled, though all caretakers and technicians were blinded to group allocation in the animal facility and to sample numbers for analysis (qPCR, titration, and histology).

#### d. Intramuscular pDNA electroporation

3B8 was delivered *in vivo*, encoded in the CMV-driven pcDNA3.4 vectors, as an equimolar mixture of the 3B8 heavy and light chain plasmids (jointly referred to as ‘p3B8’). Hamsters received an intramuscular p3B8 injection in their left and right tibialis anterior and gastrocnemius muscle (pretreated with hyaluronidase), followed by electroporation. The procedure was performed by adapting a previously optimized and validated pre-clinical protocol for mice^15^. In brief, fur at the target sites was removed using depilatory product (Veet, Reckitt Benckiser), at least two days prior to pDNA injection. Intramuscular delivery sites were injected with 100 μl of 0.4 U/μl hyaluronidase from bovine testes reconstituted in sterile saline (H4272, Sigma-Aldrich), approximately one hour prior to pDNA electrotransfer. Total p3B8 amount delivered per hamster was 600 μg (75 μl pDNA, at 2 μg/μl per muscle, formulated in sterile MQ H2O). Intramuscular injections of pDNA were immediately followed by *in situ* electroporation using the NEPA21 Electroporator (Sonidel) with CUY650P5 tweezer electrodes at a fixed width of 7 mm. Signa Electrode Gel (Parker Laboratories) was applied to the muscle to target an impedance below 0.6 Ohm. Three series of four 20 ms square-wave pulses of 120 V with a 50 ms interval were applied with polarity switching after two of the four pulses. During the procedures, hamsters were sedated using isoflurane inhalation. Electroporation parameters were based on pilot studies in hamster with a firefly luciferase reporter pDNA (data not shown). Pulse delivery was verified using the NEPA21 readout.

#### d. SARS-CoV-2 RT-qPCR

Hamster lung tissues were collected after sacrifice and were homogenized using bead disruption (Precellys) in 350 μL TRK lysis buffer (E.Z.N.A.^®^ Total RNA Kit, Omega Bio-tek) and centrifuged (10.000 rpm, 5 min) to pellet the cell debris. RNA was extracted according to the manufacturer’s instructions. RT-qPCR was performed on a LightCycler96 platform (Roche) using the iTaq Universal Probes One-Step RT-qPCR kit (BioRad) with N2 primers and probes targeting the nucleocapsid^59^. Standards of SARS-CoV-2 cDNA (IDT) were used to express viral genome copies per mg tissue^58^.

#### e. End-point virus titrations

Lung tissues were homogenized using bead disruption (Precellys) in 350 μL minimal essential medium and centrifuged (10,000 rpm, 5min, 4°C) to pellet the cell debris. To quantify infectious SARS-CoV-2 particles, endpoint titrations were performed on confluent Vero E6 cells in 96-well plates. Viral titers were calculated by the Reed and Muench method^57^ using the Lindenbach calculator and were expressed as 50% tissue culture infectious dose (TCID_50_) per mg tissue.

#### f. Sample size justification

For antiviral efficacy, we want to detect at least 1 log_10_ reduction in viral RNA levels in treated subjects compared to the untreated, infected control group. Group size was calculated based on the independent t-test with an effect size of 2.0 and a power of 80% (effect size = delta mean/SD = 1 log_10_ decrease in viral RNA/0.5 log_10_), resulting in 5-6 animals/group. Sample sizes maximized considering limits in BSL3 housing capacity, numbers of animals that can be handled under BSL3 conditions, and availability of compounds.

#### g. Ethics

Housing conditions and experimental procedures were done with the approval and under the guidelines of the ethics committee of animal experimentation of KU Leuven (license P065-2020).

### 9. Statistics

All statistical analyses were performed using GraphPad Prism 9 software (GraphPad, San Diego, CA, USA), unless indicated otherwise in the respective methods sections. Statistical significance was determined using Kruskal-Wallis test, followed by Mann Whitney U-test. P-values of <0.05 were considered significant.

## Acknowledgements

This work was funded by KU Leuven KOOR (project number ZKD8270) and KU Leuven (project SARS-CoV-2 mAb-OF2).

## Conflict of interest

The authors have no conflicts of interest.

## Author contributions.

MI, WM, KH, KV, JL, KVH, JN, PD and NG contributed to the conception and design of the work MI, WM, LA, NVdB, DVL, TV, HJT, SN, KH, XZ, DJ, RA, CF, NC, PDM, DS, JN, PD and NG contributed to the acquisition, analysis or interpretation of data

MI, WM, KH, DJ, KVH, PD and NG have drafted the work or substantively revised it

## Notes

### Competing Interest Statement

The authors have declared no competing interest.

## References

1. Ritchie, H. et al. Coronavirus Pandemic (COVID-19). https://ourworldindata.org/coronavirus (2021).

2. Walls, A. C. et al. Structure, Function, and Antigenicity of the SARS-CoV-2 Spike Glycoprotein. Cell 181, 281–292.e6 (2020).

3. Wrapp, D. et al. Cryo-EM structure of the 2019-nCoV spike in the prefusion conformation. Science (80−.). 367, 1260–1263 (2020).

4. Taylor, P. C. et al. Neutralizing monoclonal antibodies for treatment of COVID-19. Nat. Rev. Immunol. 21, 382–393 (2021).

5. Unknown. The Antibody Society: COVID-19 Biologics Tracker. https://www.antibodysociety.org/covid-19-biologics-tracker/ (2021).

6. EMA press Office,. COVID-19: EMA recommends authorisation of two monoclonal antibody medicines. 11/11/2021 https://www.ema.europa.eu/en/news/covid-19-ema-recommends-authorisation-two-monoclonal-antibody-medicines (2021).

7. Planas, D. et al. Reduced sensitivity of SARS-CoV-2 variant Delta to antibody neutralization. Nature 596, 276–280 (2021).

8. Hoffmann, M. et al. SARS-CoV-2 variants B.1.351 and P.1 escape from neutralizing antibodies. Cell 184, 2384–2393.e12 (2021).

9. Corti, D., Purcell, L. A., Snell, G. & Veesler, D. Tackling COVID-19 with neutralizing monoclonal antibodies. Cell 184, 3086–3108 (2021).

10. Ryu, D.-K. et al. Therapeutic effect of CT-P59 against SARS-CoV-2 South African variant Dong-Kyun. bioRxiv Prepr. Serv. Biol. (2021).

11. Ryu, D.-K. et al. Therapeutic efficacy of CT-P59 against P.1 variant of SARS-CoV-2. bioRxiv Prepr. Serv. Biol. (2021).

12. Ryu, D. et al. The in vitro and in vivo potency of CT-P59 against Delta and its associated variants of SARS-CoV-2 Dong-Kyun Ryu. bioRxiv Prepr. Serv. Biol. (2021).

13. Eli Lilly and Company,. FACT SHEET FOR HEALTH CARE PROVIDERS EMERGENCY USE AUTHORIZATION (EUA) OF BAMLANIVIMAB AND ETESEVIMAB. www.fda.gov (2021).

14. Hollevoet, K. & Declerck, P. J. State of play and clinical prospects of antibody gene transfer. J. Transl. Med. 15, 1–19 (2017).

15. Hollevoet, K., De Smidt, E., Geukens, N. & Declerck, P. Prolonged in vivo expression and anti-tumor response of DNAbased anti-HER2 antibodies. Oncotarget 9, 13623–13636 (2018).

16. Hollevoet, K. et al. Bridging the clinical gap for DNA-Based antibody therapy through translational studies in sheep. Hum. Gene Ther. 30, 1431–1443 (2019).

17. Vermeire, G. et al. Improved Potency and Safety of DNA-Encoded Antibody Therapeutics Through Plasmid Backbone and Expression Cassette Engineering. Hum. Gene Ther. 32, (2021).

18. CDC. SARS-CoV-2 Variant Classifications and Definitions. Sep 23 https://www.cdc.gov/coronavirus/2019-ncov/variants/variant-info.html#Interest (2021).

19. Thomson, E. C. et al. Circulating SARS-CoV-2 spike N439K variants maintain fitness while evading antibody-mediated immunity. Cell 184, 1171–1187.e20 (2021).

20. Chen, J., Gao, K., Wang, R. & Wei, G.-W. Revealing the Threat of Emerging SARS-CoV-2 Mutations to Antibody Therapies. J. Mol. Biol. 433, (2021).

21. Sanyaolu, A. et al. The emerging SARS-CoV-2 variants of concern. Ther. Adv. Vaccines 8, 1–10 (2021).

22. Lassaunière, R. et al. In vitro Characterization of Fitness and Convalescent Antibody Neutralization of SARS-CoV-2 Cluster 5 Variant Emerging in Mink at Danish Farms. Front. Microbiol. 12, 1–9 (2021).

23. Baum, A. et al. Antibody cocktail to SARS-CoV-2 spike protein prevents rapid mutational escape seen with individual antibodies. Science (80−.). 369, 1014–1018 (2020).

24. Wang, P. et al. Antibody resistance of SARS-CoV-2 variants B.1.351 and B.1.1.7. Nature 593, 130–135 (2021).

25. Zhou, D. et al. Evidence of escape of SARS-CoV-2 variant B.1.351 from natural and vaccine-induced sera. Cell 184, 2348–2361.e6 (2021).

26. CDC. COVID data tracker: variant proportions. https://covid.cdc.gov/covid-data-tracker/#variant-proportions (2021).

27. ECDC. Variants of interest and concern in the EU/EEA. https://gis.ecdc.europa.eu/portal/apps/opsdashboard/index.html#/25b6e879c076412aaa9ae7adb78d3241 (2021).

28. Abdelnabi, R. et al. Comparing infectivity and virulence of emerging SARS-CoV-2 variants in Syrian hamsters. EBioMedicine 68, 103403 (2021).

29. Food and Drug Administration. Fact Sheet for Health Care Providers Emergency Use Authorization Of REGEN-COV. (2021).

30. Connor, B. A. et al. Monoclonal Antibody Therapy in a Vaccine Breakthrough SARS-CoV-2 Hospitalized Delta (B.1.617.2) Variant Case. Int. J. Infect. Dis. 110, 232–234 (2021).

31. Hacisuleyman, E. et al. Vaccine Breakthrough Infections with SARS-CoV-2 Variants. N. Engl. J. Med. 384, 2212–2218 (2021).

32. Chen, R. E. et al. Resistance of SARS-CoV-2 variants to neutralization by monoclonal and serum-derived polyclonal antibodies. Nat. Med. 27, 717–726 (2021).

33. Collier, D. A. et al. Sensitivity of SARS-CoV-2 B.1.1.7 to mRNA vaccine-elicited antibodies. Nature 593, 136–141 (2021).

34. Kustin, T. et al. Evidence for increased breakthrough rates of SARS-CoV-2 variants of concern in BNT162b2 mRNA vaccinated individuals. medRxiv 1–17 (2021).

35. McCallum, M. et al. SARS-CoV-2 immune evasion by variant B.1.427/B.1.429. bioRxiv Prepr. Serv. Biol. (2021).

36. Wang, Z. et al. mRNA vaccine-elicited antibodies to SARS-CoV-2 and circulating variants. Nature 592, 616–622 (2021).

37. Calcoen, B. et al. Real-world monitoring of BNT162b2 vaccine-induced SARS-CoV-2 B and T cell immunity in naive healthcare workers: a prospective single center study [manuscript in preparation].

38. Tartof, S. Y. et al. Effectiveness of mRNA BNT162b2 COVID-19 vaccine up to 6 months in a large integrated health system in the USA: a retrospective cohort study. Lancet 398, 1407–1416 (2021).

39. Andrews, N. et al. Vaccine effectiveness and duration of protection of Comirnaty, Vaxzevria and Spikevax against mild and severe COVID-19 in the UK. medRxiv 2021.09.15.21263583 (2021).

40. Goldberg, Y. et al. Waning immunity of the BNT162b2 vaccine: A nationwide study from Israel. medRxiv 2021.08.24.21262423 (2021).

41. Cohn, B. A., Cirillo, P. M., Murphy, C. C., Krigbaum, N. Y. & Wallace, A. W. Breakthrough SARS-CoV-2 infections in 620,000 U.S. Veterans, February 1, 2021 to August 13, 2021. medRxiv 2021.10.13.21264966 (2021).

42. Baum, A. et al. REGN-COV2 antibodies prevent and treat SARS-CoV-2 infection in rhesus macaques and hamsters. Science (80−.). 370, 1110–1115 (2020).

43. Abdelnabi, R. et al. Comparative infectivity and pathogenesis of emerging SARS-CoV-2 variants in Syrian hamsters. EBioMedicine 68, 1–26 (2021).

44. Andreano, E. et al. Extremely potent human monoclonal antibodies from COVID-19 convalescent patients. Cell 184, 1821–1835.e16 (2021).

45. Kreye, J. et al. A Therapeutic Non-self-reactive SARS-CoV-2 Antibody Protects from Lung Pathology in a COVID-19 Hamster Model. Cell 183, 1058–1069.e19 (2020).

46. Fagre, A. C. et al. A Potent SARS-CoV-2 Neutralizing Human Monoclonal Antibody That Reduces Viral Burden and Disease Severity in Syrian Hamsters. Front. Immunol. 11, 1–13 (2020).

47. Piepenbrink, M. S. et al. Therapeutic activity of an inhaled potent SARS-CoV-2 neutralizing human monoclonal antibody in hamsters. Cell Reports Med. 2, 100218 (2021).

48. Winkler, E. S. et al. Human neutralizing antibodies against SARS-CoV-2 require intact Fc effector functions for optimal therapeutic protection. Cell 184, 1804–1820.e16 (2021).

49. Tortorici, M. A. et al. Ultrapotent human antibodies protect against SARS-CoV-2 challenge via multiple mechanisms. Science (80−.). 370, 950–957 (2020).

50. Hansen, J. et al. Studies in humanized mice and convalescent humans yield a SARS-CoV-2 antibody cocktail. Science (80−.). 369, 1010–1014 (2020).

51. Jones, B. E. et al. LY-CoV555, a rapidly isolated potent neutralizing antibody, provides protection in a non-human primate model of SARS-CoV-2 infection. bioRxiv Prepr. Serv. Biol. 1–29 (2020) doi:10.1101/2020.09.30.318972.

52. Andrews, C. D., Huang, Y., Ho, D. D. & Liberatore, R. A. In vivo expressed biologics for infectious disease prophylaxis: rapid delivery of DNA-based antiviral antibodies. Emerg. Microbes Infect. 9, 1523–1533 (2020).

53. Lee, W. S., Wheatley, A. K., Kent, S. J. & DeKosky, B. J. Antibody-dependent enhancement and SARS-CoV-2 vaccines and therapies. Nat. Microbiol. 5, 1185–1191 (2020).

54. Picelli, S. et al. Full-length RNA-seq from single cells using Smart-seq2. Nat. Protoc. 9, 171–181 (2014).

55. Ozawa, T., Kishi, H. & Muraguchi, A. Amplification and analysis of cDNA generated from a single cell by 5’-RACE: application to isolation of antibody heavy and light chain variable gene sequences from single B cells. Biotechniques 40, 469–478 (2006).

56. Sanchez-Felipe, L. et al. A single-dose live-attenuated YF17D-vectored SARS-CoV-2 vaccine candidate. Nature 590, 320–325 (2021).

57. Reed, L. & Muench, H. A simple method of estimating fifty per cent endpoints. Am J Epidemiol. 27, 493–497 (1938).

58. Kaptein, S. J. F. et al. Favipiravir at high doses has potent antiviral activity in SARS-CoV-2-infected hamsters, whereas hydroxychloroquine lacks activity. Proc. Natl. Acad. Sci. U. S. A. 117, 26955–26965 (2020).

59. Boudewijns, R. et al. STAT2 signaling restricts viral dissemination but drives severe pneumonia in SARS-CoV-2 infected hamsters. Nat. Commun. 11, 1–10 (2020).

