## Supplemental data for "Potent neutralizing anti-SARS-CoV-2 human antibodies cure infection with SARS-CoV-2 variants in hamster model"

### Supplementary data

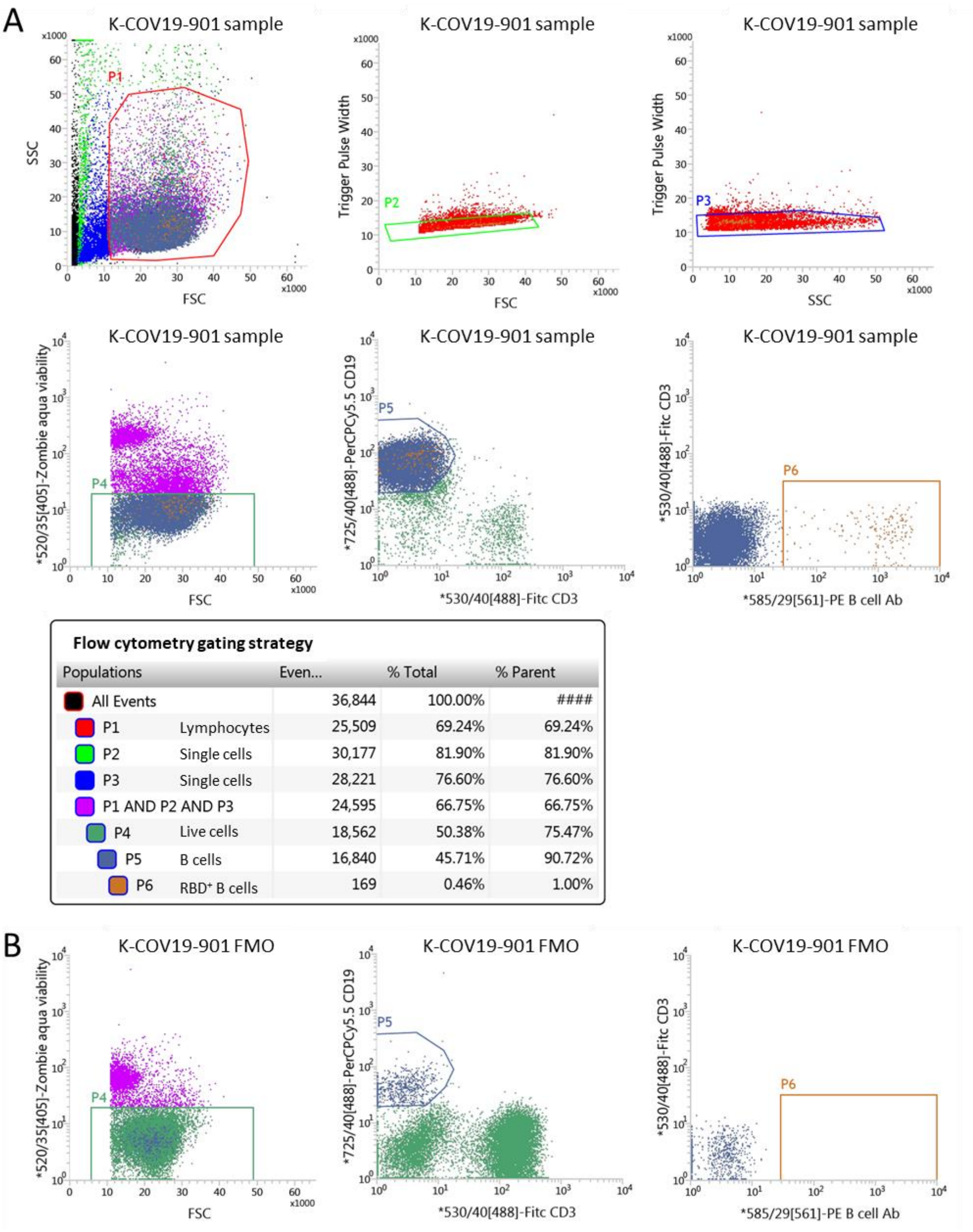

**Flow cytometry gating strategy**

| Populations | | Even... | % Total | % Parent |
|---|---|---|---|---|
| All Events | | 36,844 | 100.00% | #### |
| P1 | Lymphocytes | 25,509 | 69.24% | 69.24% |
| P2 | Single cells | 30,177 | 81.90% | 81.90% |
| P3 | Single cells | 28,221 | 76.60% | 76.60% |
| P1 AND P2 AND P3 | | 24,595 | 66.75% | 66.75% |
| P4 | Live cells | 18,562 | 50.38% | 75.47% |
| P5 | B cells | 16,840 | 45.71% | 90.72% |
| P6 | RBD+ B cells | 169 | 0.46% | 1.00% |

Supplementary figure 1: gating strategy for selection of RBD-specific B cells from PBMCs of patients K-COV19-901. Lymphocytes were selected based on FSC and SSC, followed by selection of single cells and live cells (Zombia Aqua negative). Here, B cells were selected as CD19+ CD3- cells and evaluated for RBD surface staining using PE as fluorophore. Fully stained sample is shown in panel A, fluorescence minus one (FMO) control for RBD fluorescence is shown in panel B.

Supplementary Table 1: overview of RBD-binding titers and neutralizing antibody titers in serum, and percentage RBD-positive B cells in PBMCs from 26 convalescent COVID-19 patients

| **Patient ID** | **anti-RBD Ab (µg/ml)** | **Neutralizing Abs (SNT50 titer)** | **% RBD-positive of B cells** |
|---|---|---|---|
| K-COV19-901 | 182,16 | 1209 | 0,80% |
| K-COV19-902 | 21,65 | 96 | 0,00% |
| K-COV19-006 | 387,59 | 7022 | 0,00% |
| K-COV19-007 | 5,98 | / | 0,00% |
| K-COV19-009 | 7,57 | 8 | 0,04% |
| K-COV19-010 | 15,58 | 74 | 0,03% |
| K-COV19-019 | 26,07 | 771 | 0,05% |
| K-COV19-021 | 30,80 | 506 | 0,19% |
| K-COV19-028 | 25,75 | 493 | 0,00% |
| K-COV19-034 | 48,82 | 810 | 0,02% |
| K-COV19-041 | 8,66 | 201 | 0,09% |
| K-COV19-044 | 40,22 | 876 | 0,00% |
| L-COV19-001 | 14,75 | 178 | 0,14% |
| L-COV19-002 | 24,37 | 248 | 0,30% |
| L-COV19-003 | 51,37 | 481 | 0,18% |
| L-COV19-004 | 48,71 | 302 | 0,07% |
| L-COV19-005 | 196,52 | 2123 | 0,14% |
| L-COV19-007 | 32,00 | 829 | 0,22% |
| L-COV19-008 | 54,97 | 1539 | 0,35% |
| L-COV19-009 | 32,07 | 1199 | 0,21% |
| L-COV19-010 | 16,76 | 104 | 0,19% |
| L-COV19-011 | 32,26 | 495 | 0,11% |
| L-COV19-012 | 13,99 | 140 | 0,07% |
| L-COV19-015 | 41,20 | 3077 | 0,14% |
| L-COV19-018 | 10,36 | 69 | 0,11% |

Supplementary Table 2: overview of single RBD mutants used in this study and their presence in different SARS-CoV-2 variants of concern, variants of interest, variants under monitoring or de-escalated variants. Information retrieved from European Centre for Disease Prevention and Control (https://www.ecdc.europa.eu/en/covid-19/variants-concern)

| | **SARS-CoV-2 variants** | | | | |
|---|---|---|---|---|---|
| **Mutation** | **Concern** | **Interest** | **Monitored** | **De-escalated*** | **Other** |
| **N439K** | | | | AV.1 | Ab evasion mutant |
| **L452R** | B.1.617.2 (delta) | C.37 (Lambda) | AY.4.2; | B.1.427/B.1.429 (Epsilon); B.1.617.1 (Kappa); B.1.617.3; A.27; C.16; B.1.526.1 | |
| **Y453F** | | | | | Mink mutant, Ab evasion mutant |

| | | | | |
|---|---|---|---|---|
| **E484K** | B.1.351 (beta); P.1 (Gamma) | B.1.621 (Mu) | B.1.1.318; C.1.2 | B.1.525 (Eta); P.3 (Theta); B.1.620; A.28; B.1.526 (Iota); P.2 (Zeta); AT.1; AV.1 |
| **E484Q** | | | | B.1.617.1 (Kappa); B.1.617.3 |
| **N501Y** | B.1.351 (beta); P.1 (Gamma) | B.1.621 (Mu) | | B.1.1.7 (Alpha); P.3 (Theta); A.27 |

**These variants of SARS-CoV-2 have been de-escalated based on at least one the following criteria: (1) the variant is no longer circulating, (2) the variant has been circulating for a long time without any impact on the overall epidemiological situation, (3) scientific evidence demonstrates that the variant is not associated with any concerning properties (https://www.ecdc.europa.eu/en/covid-19/variants-concern).*
